# Associative synaptic plasticity creates dynamic persistent activity

**DOI:** 10.1101/2025.08.20.671131

**Authors:** Albert J. Wakhloo, David G. Clark, L.F. Abbott

**Affiliations:** Center for Theoretical Neuroscience, Zuckerman Institute, Columbia University; Center for Computational Neuroscience, Flatiron Institute, New York, NY, 10010; Kavli Institute for Brain Science, Columbia University, New York, NY, 10027; Kempner Institute for the Study of Natural and Artificial Intelligence, Harvard University, Boston, MA, 02134

**Keywords:** cellular automata, Wolfram rules, finite fields, reaction–diffusion, Turing patterns, graph Laplacian, cochain complex, cohomology, morphogenesis

## Abstract

In biological neural circuits, the dynamics of neurons and synapses are tightly coupled. We study the consequences of this coupling and show that it enables a novel form of working memory. In recurrent neural network models with ongoing Hebbian plasticity, we find that following oscillatory stimulation, neurons continue to oscillate long after the input is removed. This creates a dynamic form of memory that has no explicit storage or retrieval phases and that requires no prior knowledge of the input. We trace the mechanism of these “persistent oscillations” to an interaction between neurons and synapses that creates complex outlier eigenvalues of the connectivity matrix. This is shown both in simulation and analytically. We leverage this mechanistic understanding to generate persistent oscillations with prespecified dynamics, creating a dynamic analog of a classical Hopfield network. Our work demonstrates that coupling neuronal and synaptic dynamics enables novel forms of computation.

## I. INTRODUCTION

Synaptic connections shape neuronal dynamics, and neuronal dynamics modify synapses through activity-dependent plasticity mechanisms. The dynamics of neurons and synapses are therefore tightly coupled. Nevertheless, classical neural-network models treat these processes separately, with models of neuronal dynamics typically assuming fixed synapses [1–5] and models of learning and memory typically separating plasticity from the dynamics of the network [6–9]. Most efforts to study the computational functions of joint neuronal-synaptic dynamics have limited their focus to purely presynaptic short-term plasticity [10–14]. However, experimental work has described forms of bidirectional, associative plasticity over timescales not that different from those of purely neural systems [15–19]. Thus, it is interesting to ask what unique network functions are enabled by ongoing associative plasticity [20–26].

In this paper, we study how recurrent neural networks with ongoing associative plasticity respond to external stimulation. In this model, introduced by Clark and Abbott [25], the synapses of a recurrent neural network evolve about random baseline strengths according to a Hebbian rule (Fig. 1A), and neurons and synapses fluctuate together on comparable timescales. Importantly, plasticity influences neuronal dynamics not by creating entirely new dynamical motifs that erase the prior weights, but by interacting with and sculpting a baseline connectivity. This requires the size of the synaptic fluctuations to be small–i.e., individual synapses remain close to their baseline strengths (Fig. 1B-C).

**FIG. 1.**
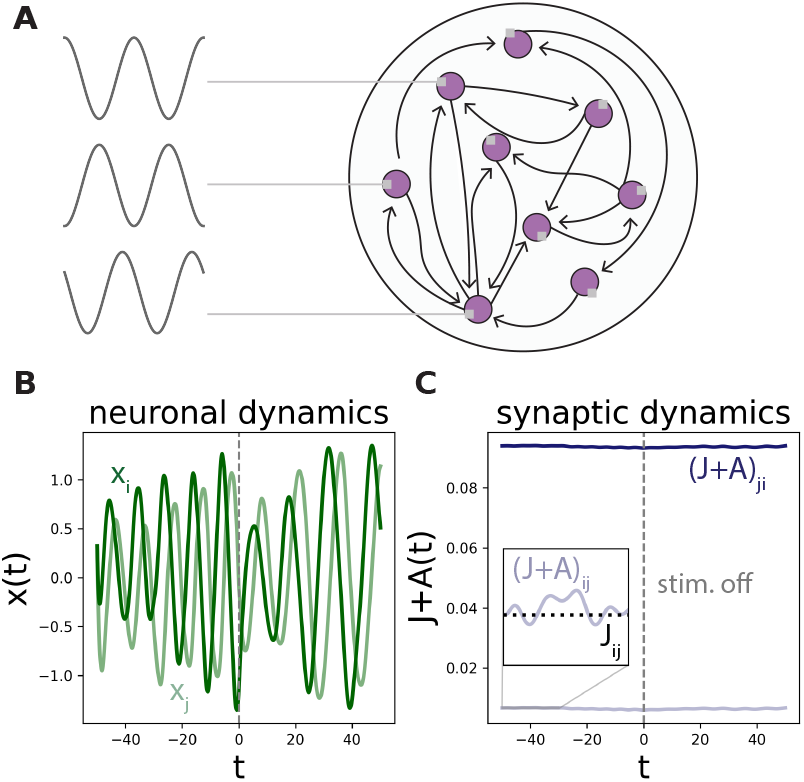
(A) Schematic of our modeling framework. (B) Neuronal dynamics of two example units, *x*_*i*_ and *x*_*j*_, from a network of *N* = 500 units during the stimulation period (*t <* 0) and following input cessation (*t >* 0). Here and in the remainder of the paper, we measure time relative to the moment the input is turned off (grey dashed line) and in units of the neuronal time constant of the network. (C) Dynamics of the corresponding synaptic weights between these two units, *J*_*ij*_ + *A*_*ij*_ (*t*) and *J*_*ji*_ + *A*_*ji*_(*t*). Fluctuations in the synaptic weights (*J*_*ij*_ + *A*_*ij*_ (*t*)) (inset, gray line) are small relative to the baseline weights (*J*_*ij*_ ) (inset, dashed line).

We study the response of such networks to a prototypical class of inputs, oscillations [27]. Our main finding is that the interaction of recurrent neural dynamics with Hebbian synaptic dynamics creates oscillatory activity that outlasts the external stimulation for a time much longer than the intrinsic timescales of the system. This is unexpected, because Hebbian plasticity typically leads to fixed points, as in Hopfield models [6]. We show that Hebbian associative plasticity rules can generate these more complex dynamical patterns as a result of the interaction between the static backbone connectivity in our model and ongoing plasticity.

## II. RESULTS

### A. Recurrent-network model with ongoing Hebbian plasticity and external drive

We study the recurrent-network model of Clark and Abbott [25] in which synapses fluctuate around fixed, random baselines. The dynamic variables comprise *N* neurons, with preactivations denoted by *x*_*i*_(*t*), together with *N* ^2^ plastic synaptic weights *A*_*ij*_(*t*), where *i, j* range from 1 to *N* . The neuronal variables evolve according to

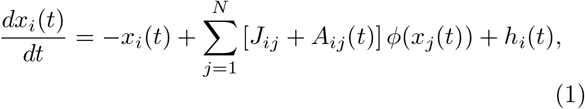

where the variables *J*_*ij*_ are fixed random weights drawn from a Gaussian with mean zero and standard deviation 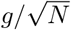 (although our results are expected to apply for non-Gaussian weight distributions as well). These weights form the static backbone of the connectivity, while the dynamic variables *A*_*ij*_(*t*) are fluctuations due to ongoing plasticity. These are denoted in matrix form by ***J*** and ***A***(*t*). The function *ϕ*(·) is a saturating nonlinearity mapping preactivations to shifted and normalized firing rates, which we set to a hyperbolic tangent, *ϕ*(·) = tanh(·). In addition to the recurrent input from other units, each unit receives an external input *h*_*i*_(*t*), defined below.

The time-varying component of the connectivity evolves according to a “pre-times-post” rate-based Hebbian rule,

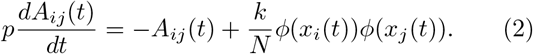

Here, *p* sets the timescale of the synaptic dynamics relative to the neuronal dynamics, and *k* determines the strength and sign of the plasticity. We consider *k* ≥ 0, corresponding to a Hebbian (rather than anti-Hebbian) rule, and *p >* 1, so that synapses evolve more slowly than neurons. Importantly, while the static background connections, *J*_*ij*_, scale as 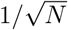, the plasticity-induced changes to these connections, *A*_*ij*_(*t*), scale as 1*/N* . As we focus on the large-*N* regime, the plastic component is vanishingly small relative to the static component. Such minuscule changes in the synapses can, nevertheless, have dramatic effects on the network dynamics due to network activity aligning to low-rank structure present in ***A***(*t*) [25].

In contrast to Clark and Abbott [25], we introduce a nonzero external input *h*_*i*_(*t*) given by

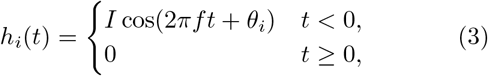

with amplitude *I*, frequency *f*, and neuron-specific phase *θ*_*i*_. We choose the phases randomly for each unit following Rajan et al. [27], though our results are qualitatively unchanged when all phases are equal (*θ*_*i*_ = *θ*).

### B. Behavior of the externally driven, plastic network

We first study the behavior of this model when the input is active, *t <* 0. This extends the phenomenology studied by Rajan et al. [27], which considered static synapses, to the model of Clark and Abbott [25], which includes plastic synapses. Neuronal traces are visualized in Fig. 2Ai–iv (*t <* 0). When both the input amplitude, *I*, and plasticity strength, *k*, are small, neurons show chaotic fluctuations with superimposed oscillations (Fig. 2Ai). As the plasticity strength *k* increases, the chaotic activity progressively slows (Fig. 2Aii). This occurs because plastic synapses create a positive feedback loop that slows the neuronal dynamics and drags the network toward fixed points [25]. In this state, external input produces small oscillations atop the nearly static activity. When the input amplitude *I* increases sufficiently, the network transitions from this “small-oscillation” regime to a state that contains both a stimulus-driven component and, due to the recurrent connectivity, an intrinsically generated chaotic component (Fig. 2Aiii). As *I* continues to increase, the neurons eventually become perfectly entrained to the input, eliminating all traces of chaos. (Fig. 2Aiv, Methods).

**FIG. 2.**
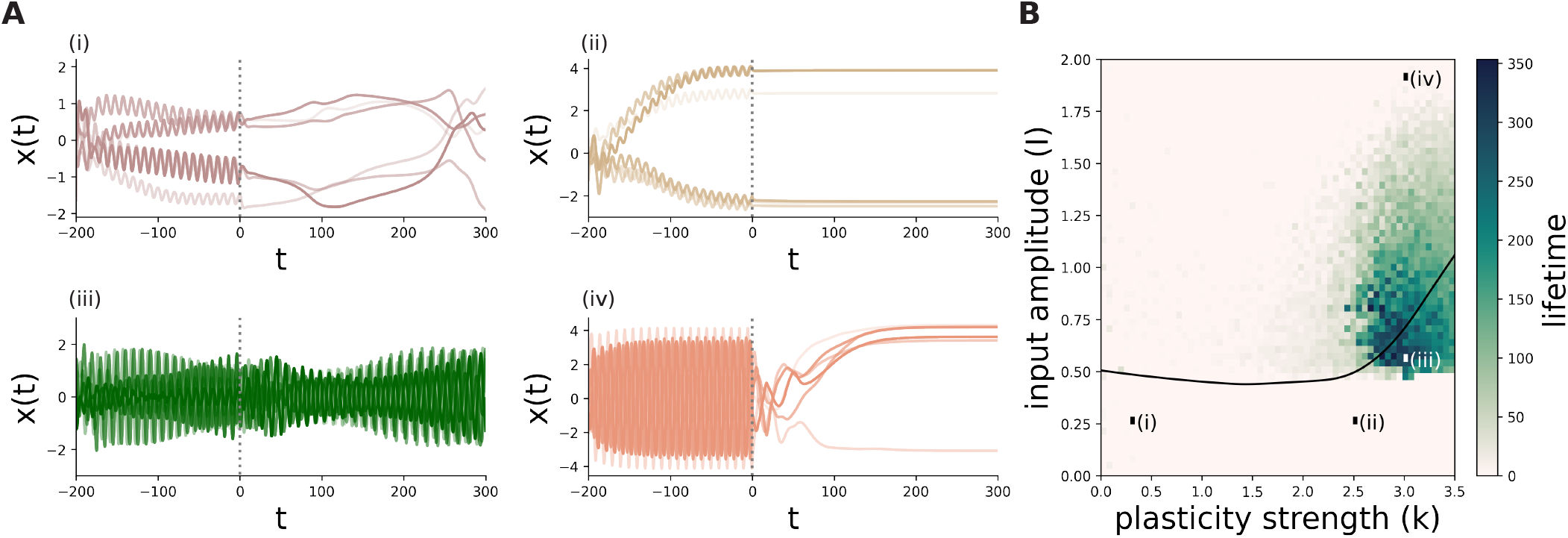
Overview of model behavior. (A) Example traces at four different points of the parameter space. In each subplot, we stimulate a network for 200 time units, after which point all inputs are halted. The duration of the stimulation is not essential, provided it is sufficiently long (SM Fig. 2). Each line denotes a single unit from the same network. (B) Lifetime of persistent oscillations as a function of *k* and *I* for networks with *p* = 25, *g* = 1.3, and *f* = 0.1. The black line marks the transition from chaos to perfectly oscillatory activity during stimulation (Methods). Each point in the (*k, I*) plane corresponds to an average over 50 networks of size *N* = 500.

### C. Persistent oscillations following input cessation

The four regimes shown in (Fig. 2A) exhibit distinct behaviors following input cessation at *t* = 0 (Fig. 2Ai–iv, *t* ≥ 0). Most notably, the network can continue to oscillate after the input is removed (Fig. 2Aiii). The lifetime of these oscillations can exceed the synaptic timescale, *p*, by an order of magnitude (Fig. 2B and Fig. 3A, Methods). The fact that these oscillations persist for much longer than the intrinsic timescales of the system indicates that they reflect a *self-renewing process*. We demonstrate this in SM Fig. 1by setting *k* = 0 at *t* = 0 so that synapses continue to influence neurons, and continue to decay, but neurons no longer influence synapses. In this case, persistent oscillations are curtailed. These results show that persistent oscillations involve oscillatory neural activity “feeding back into itself” through ongoing synaptic weight changes.

**FIG. 3.**
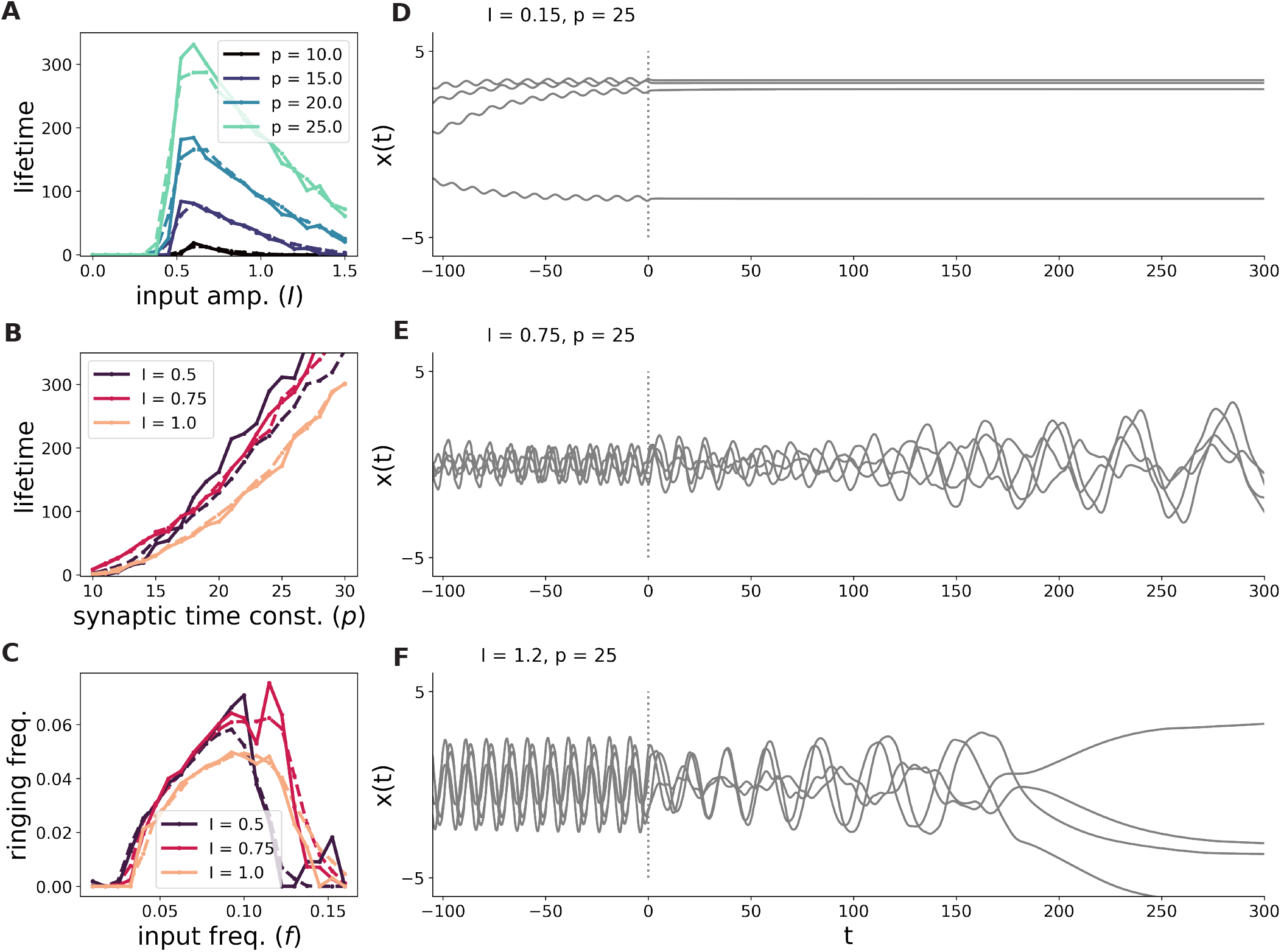
Scaling of lifetime and frequency with model parameters. (A-B) Lifetime of persistent oscillations as a function of input amplitude *I* and synaptic time constant *p* for networks with *N* = 1, 000 (dashed) and *N* = 5, 000 (solid). Each point represents an average over 400 simulations for the small networks and over 30 for the large networks. (C) Frequency of the initial phase of the persistent oscillation as a function of the frequency of the inputs. We define this “ringing frequency” by the first peak in the power spectrum of the neuronal dynamics (Methods). When no peak is detected, we set the frequency to 0. (D-F) In each subplot, we plot the activity of four example units from three different networks with increasing values of *I* and with all other parameters held fixed (first 100 time units of stimulation not shown). When *I* is too low (panel D; *I* = 0.15), the network is in the small oscillation regime. When it is moderately large (E; *I* = 0.75), the network expresses a mixture of recurrent- and input-driven features, and oscillations persist for a long time. Finally, as *I* continues to increase (F; *I* = 1.2), the activity becomes increasingly dominated by the inputs, and the lifetime of the oscillations gradually decreases.

We now describe the dependence of persistent oscillations on system parameters. Here, and for the rest of the paper, we focus on the regime in which synapses are slower than neurons (*p >* 1) and where, without plasticity, the activity would be chaotic (*g >* 1) [1, 25]. As shown in Fig. 3B, the lifetime of persistent oscillations grows supralinearly with *p*. For these oscillations to occur, the input amplitude, *I*, must be large enough to avoid the small-oscillation regime, but small enough to avoid eliminating the influence of the recurrent connections on the dynamics (Figs. 2Aii and 3A). That is, the activity must express a mixture of inputand recurrentdriven features for persistent oscillations to occur.

The fact that persistent oscillations are eliminated for large *I* shows that this phenomenon is not the result of a simple delayed feedback loop between neurons and synapses. Indeed, one would expect the opposite trend if this were the mechanism driving persistent oscillations. We also note that the parameter regimes in which persistent oscillations do or do not occur are distinct from those defining the phases of chaos or oscillatory entrainment described in [27] (Fig. 2B).[28]

We next consider the relationship between the frequency of the input and the frequency of persistent oscillations (Methods). As shown in Fig. 3C, there is a preferred frequency band within which oscillations persist. Inside this band, the frequency of the oscillations grows with the frequency of the input. As the input frequency approaches the upper edge of the band, the likelihood of generating persistent oscillations rapidly falls. These findings imply that when oscillations persist, they reflect the structure of the previously present inputs, with faster inputs leading to faster persistent oscillations.

We expect these trends in the lifetime and frequency of persistent oscillations to hold in the *N* → ∞ limit. While it is possible to derive a dynamical mean field theory that describes this limit exactly [25], we found that the resulting equations were prohibitively difficult to solve numerically. Instead, we show in Fig. 3 that all trends discussed above are identical in networks with *N* = 1000 and *N* = 5000.

### D. Complex outlier eigenvalues underlie persistent oscillations

How does the connectivity matrix generate oscillations in the absence of inputs? We begin by considering the eigenvalues of the matrix ***J*** + ***A***(*t*) at the time the input is turned off, *t* = 0. We expect these eigenvalues to be predictive of the nonlinear dynamics of the system, with large, complex eigenvalues leading to oscillations.

This is indeed the case. In the parameter regime where oscillations persist, the connectivity matrix develops a pair of complex-conjugate outlier eigenvalues (Fig. 4Ai, outliers). The distribution of the remaining eigenvalues is a uniform disk in the complex plane of radius *g*, which is the eigenvalue distribution of an i.i.d. random matrix—i.e. the eigenvalue distribution for ***J*** alone, without plastic modifications (Fig. 4(A-B)i, disk). In the small-oscillation regime in which oscillations do not persist, the connectivity develops a single large real outlier eigenvalue, reflecting the neurons being continuously dragged towards fixed points by the synapses (Fig. 4Bi).

**FIG. 4.**
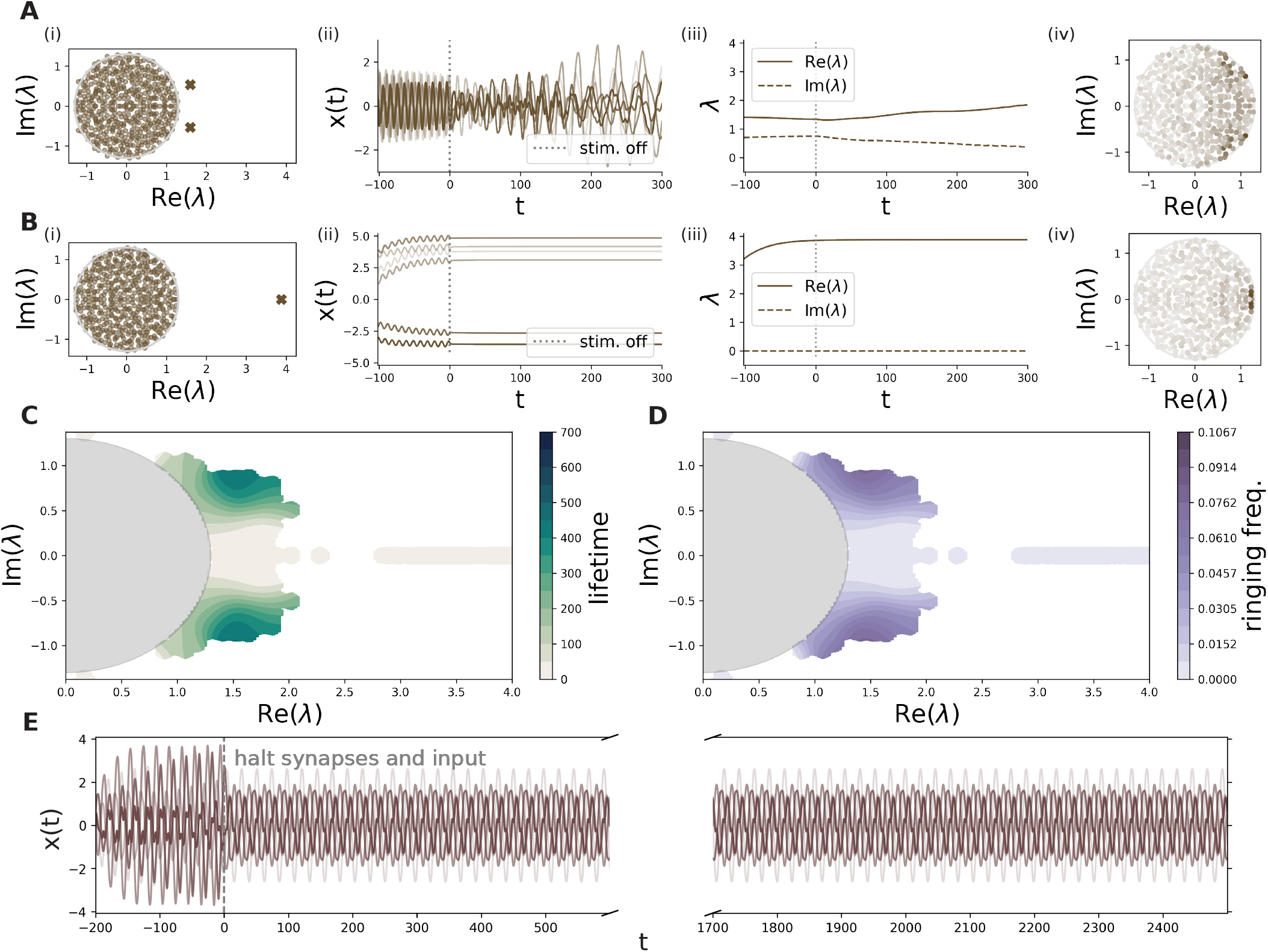
Eigenvalue outliers drive persistent oscillations. (A-B) Behavior of outlier eigenvalues when the network is in the persistent oscillation regime (A), versus when it is in the small oscillation/slow dynamics regime (B). (i) Eigenvalues of ***J*** +***A***(*t*) at *t* = 0, when the input is halted. The bulk of the eigenvalue distribution fills a disk of radius *g* (gray line), which is what one would expect from a random matrix with variance *g*^2^*/N*, while the outliers are produced by the plastic matrix. (ii) Dynamics of 5 example units, *x*_*i*_(*t*), from the corresponding network (first 100 time units not shown). (iii) Dynamic trajectory of the real and imaginary parts of the largest outlier eigenvalue of the network. (iv) Overlaps of the primary eigenvectors of the plastic ***A***(*t*) matrix with each eigenvector of ***J*** at the time the input is halted. Here we color each eigenvalue of ***J*** by the magnitude of the corresponding eigenvector’s alignment with ***A***(0) (Methods). In the persistent oscillation regime, the plastic weights align with eigenvectors of ***J*** with complex eigenvalues. We return to this point in Sec. II F. (C-D) The lifetime and frequency of persistent oscillations as a function of the eigenvalue outlier, with the expected disk-shaped eigenvalue bulk shaded in gray. We simulated a set of networks over a range of parameters and plotted the resulting eigenvalue outliers, coloring by either lifetime or persistent oscillation frequency (D) and binning the final values (Methods). (E) Freezable persistent oscillations. Here, we halt the ongoing plasticity at *t* = 0, creating persistent oscillations that never decay. This is a dynamic variant of freezable chaos–see Ref. [25].

Of course, the connectivity is actually dynamic, and so too are the outlier eigenvalues. The dynamics of the outlier eigenvalues reflect the behavior of the network after the input is turned off. As oscillations persist (Fig. 4Aii), the imaginary component of the eigenvalues gradually falls to zero while the real part grows more positive (Fig. 4Aiii). During this time, the oscillations progressively slow, eventually stopping altogether around the time the outlier becomes real. In SM Fig. 5, we show that the instantaneous network frequency closely tracks the imaginary component of the eigenvalue outlier through the entire period following input cessation, providing further evidence that outliers drive persistent oscillations.

**FIG. 5.**
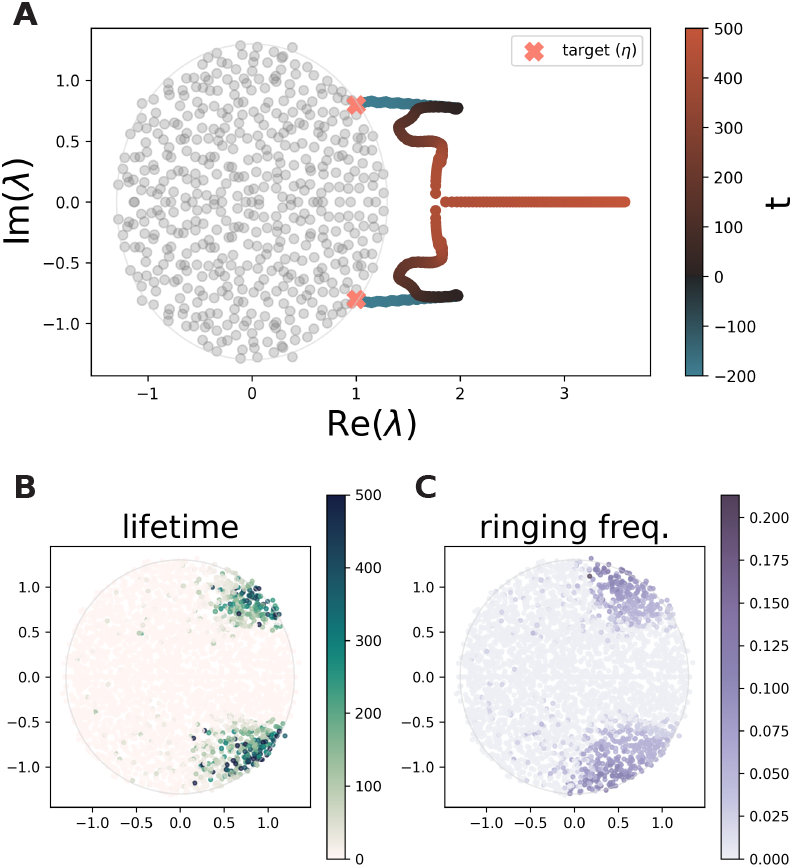
Generating structured persistent oscillations with targeted inputs. (A) Trajectory of a targeted eigenvalue. Here, we target an eigenvalue located at 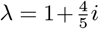 and track its trajectory during the stimulation period (blue) and the stimulus off period (red). As in previous figures, *t* denotes the time from the moment the stimulation is halted. (B-C) Lifetime and ringing frequency as a function of the targeted eigen-value. The real part affects the lifetime, while the imaginary part affects both the lifetime and the frequency of the evoked oscillation.

The locations of the outlier eigenvalues largely determine the lifetime and frequency of the persistent oscillations. To show this, we sweep over a large number of model parameters and plot the lifetime and frequency of the oscillations as a function of the real and imaginary parts of the outlier at the time the input is turned off (Fig. 4C-D). In general, oscillations have a longer lifetime when the outlier has a moderately large real part and have both a higher frequency and a longer lifetime when the outlier has a large imaginary part. That is, the imaginary component affects both the lifetime and the frequency of the oscillations, while the real part controls only the lifetime.

We now use this mechanism to generate persistent oscillations that never decay. To do this, we freeze all synaptic dynamics at *t* = 0, halting both Hebbian updates and synaptic decay (effectively sending *p* → ∞ as in [25]). This fixes the synaptic weights and eigenvalue spectrum at their post-stimulation values. With plasticity frozen, oscillations induced by the outlier eigenvalues persist indefinitely (Fig. 4E).

The emergence of complex-conjugate outlier eigenvalues is surprising given that the Hebbian rule produces symmetric updates to the weights (***A***(*t*) = ***A***^⊤^(*t*); Eq. 2), and symmetric matrices have purely *real* eigenvalues. Moreover, for any low-rank, symmetric matrix ***H*** = ***H***^⊤^ that does not depend on ***J***, the combined matrix ***J*** + ***H*** can possess only real outlier eigenvalues in the large-*N* limit (SM Sec. 2.1; note that, in our model, the plastic matrix ***A***(*t*) is approximately low-rank [25] due to *p* being finite) [29]. These outliers, if they exist, are simply the (real) eigenvalues of ***H*** (SM Sec. 2.1).

Thus, for *complex* outliers to emerge, the plastic matrix cannot be uncorrelated with the static background matrix ***J*** . Because ***A***(*t*) is shaped by activity-dependent plasticity, complex outliers and persistent oscillations can only occur when the activity expresses features driven by both the external inputs and the recurrent background connectivity, ***J*** . We visualize this in Fig. 4Aiv, which demonstrates that in the persistent oscillation regime, ***A***(*t*) is aligned with the subspaces of eigenvectors of ***J*** corresponding to complex eigenvalues (Methods).

In the following section, we demonstrate this idea by first showing that a matrix built from eigenvectors of ***J*** is sufficient for producing complex outliers and thus oscillations, and then designing inputs that generate such a matrix via Hebbian plasticity during the stimulation period.

### E. Generating targeted persistent oscillations

We now address how the interaction between the static backbone, ongoing plasticity, and oscillatory inputs can generate complex outlier eigenvalues in the connectivity matrix. We first define a “target” matrix ***Â*** in terms of eigenvectors of ***J*** that, when added to ***J***, generates complex outlier eigenvalues and thus oscillations. We then find a set of inputs that allow Hebbian plasticity to generate such a matrix during the stimulation period, that is, stimulation leading to ***A***(0) ≈ ***Â***. This approach is similar to that of Hopfield networks in which targeted inputs cause a network to express a pattern of activity embedded in the synapses. Here, however, the memory-related activity is dynamic, rather than static, and memory storage is part of the dynamics of the model.

The target matrix is rank-two, real, and symmetric, given by

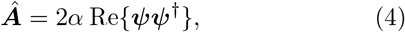

where ***ψ*** is a complex eigenvector of ***J***, † denotes the complex conjugate transpose, and *α* is a real scalar. We take ***ψ*** to be a right eigenvector of ***J***, that is, ***Jψ*** = *η****ψ***, with eigenvalue *η*. In SM Sec. 2.2, we show that for any choice of ***J*** (i.e., not necessarily random), the combined matrix ***J*** + ***Â*** develops an eigenvalue pair located at

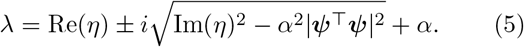

In the large-*N* limit with ***J*** i.i.d. Gaussian, |***ψ***^⊤^***ψ***|^2^ ≪ 1, so that the above equation simplifies to

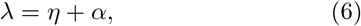

together with its conjugate. In other words, the matrix ***Â*** translates the eigenvalue *η* along the real axis; this translation is rightward for *α >* 0.

Furthermore, we provide a stimulation protocol to create a plastic matrix similar to ***Â*** above. If the neuronal activations were perfectly sinusoidal with *ϕ*(***x***(*t*)) = Re{***ψ****e*^*i*2*πft*^} and *p* ≫ 1*/f*, Eq. 2 shows that the plastic matrix ***A***(*t*) would converge to the ***Â*** of Eq. 4 with *α* = *k/*4*N* . Of course, the actual neuronal variables are nonlinear, and it is in general not possible to force them to be exact sinusoids. Nevertheless, we can stimulate the network with such perfectly sinusoidal inputs given by

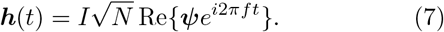

The factor of 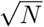 ensures that these inputs are of order one for the conventional normalization ∑_*i*_ |*ψ*_*i*_|^2^ = 1 . These sinusoidal inputs have phases that are aligned with the chosen eigenvectors of the static connectivity matrix, ***J*** .

As shown in Fig. 5A, this eigenvector-aligned input selectively amplifies specific complex-conjugate eigenvalues, translating them rightward along the real axis and creating a complex outlier pair that drives persistent oscillations. Figs. 5B and 5C illustrate the relationship between the lifetime and frequency of the persistent oscillations and the target eigenvalue *η*. The lifetime is controlled by both the real and imaginary parts of *η*, while the frequency is controlled by its imaginary part alone. Using this eigenvalue-targeting method, we can generate persistent oscillatory activity with controlled lifetime and frequency characteristics.

The relationship between system parameters and persistent oscillations is different in this eigenvector-aligned case than in the random-phase case explored previously. Most notably, with eigenvector-aligned inputs, we can use arbitrarily large input amplitudes and still elicit persistent oscillations. This key difference arises because the stimulus is now explicitly correlated with the random recurrent connectivity. Consequently, even when network activity is input-dominated, the plastic matrix ***A***(*t*) still develops correlations with the random backbone, ***J*** .

We can show analytically that, with eigenvector-aligned inputs and taking *p, I*, and 1*/f* large with *p* ≫ 1*/f*, the resulting eigenvalue outliers following a sufficiently long stimulation period are approximated by

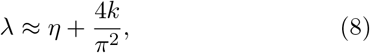

and its conjugate 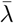 (SM Sec. 3.1), if |*λ*| *> g* (otherwise no outliers exist). The coefficient 4*/π*^2^ results from the effects of the nonlinearity *ϕ*(·). This expression indicates that the eigenvalue outlier—and thus the lifetime and frequency of the targeted oscillation—depends only on the plasticity strength *k* and the targeted eigenvalue *η*. Eigenvector-aligned inputs therefore enable persistent oscillations with arbitrarily large input amplitudes *I*, while also making these oscillations insensitive to the exact input frequency *f* . Instead, the oscillations depend on the targeted connectivity mode encoded in the pattern of input weights across neurons.

These results demonstrate that targeted inputs decouple stimulus-evoked dynamics from the persistent dynamics that follow. What matters is the “spatial” pattern of the inputs across neurons (i.e., their alignment with a particular eigenvector), not their temporal structure. We show an extreme example of this in SM Fig. 4, where we evoke persistent oscillations with a single biphasic pulse rather than sustained oscillatory drive. This behavior differs from the random-phase case, in which input amplitude and frequency affect the persistent oscillations that follow (Fig. 3).

In the next section, we analyze the random-phase case and show that it too involves eigenvector alignment, unifying the two cases. We note, however, that eigenvector alignment may not be the only route to complex outliers; other low-rank perturbations of ***J*** could produce them as well, a possibility we leave to future work.

### F. Analytical solution for random phases: outlier location and eigenvector alignment

How do complex outlier eigenvalues emerge in response to random-phase oscillatory input (Fig. 4), and is this mechanism related to the eigenvector alignment for targeted inputs described above? To answer these questions, we obtain an analytical approximation for the plastic matrix ***A***(0) and its resulting outlier eigenvalues as a function of the model parameters. This approximation reveals that the random-phase case also involves alignment between ***A***(0) and the eigenvectors of ***J***, unifying the two cases.

To motivate our analysis, note that the key features of the dynamics during stimulation are that *x*_*i*_(*t*) is 1) driven by the input *I*_*i*_(*t*), and 2) also influenced by ***J*** (Fig. 2). We reasoned that these features could be captured by a linearized model in which we make the approximation *ϕ*(*x*_*i*_(*t*)) ≈ *γx*_*i*_(*t*), with a gain parameter *γ* that is set heuristically (SM Sec. 3.2).

As described in SM Sec. 3.2, this approach yields the following approximation for the dynamics of the activations:

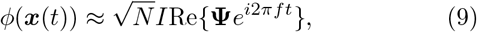

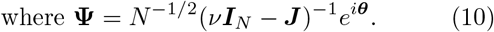

Here, *e*^*i****θ***^ is a vector with elements 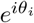 with *θ*_*i*_ being the phase of the input to neuron *i*, ***I***_*N*_ is the identity matrix, and *ν* is a complex scalar that depends on system parameters (see Methods for the full expression).

Expanding the above equation in powers of ***J***, we find that the parameter *ν* measures the degree to which the neuronal dynamics are influenced by the recurrent connectivity. For large values of |*ν*|, the variables *ϕ*(*x*_*i*_(*t*)) are dominated by the inputs, while for |*ν*| close to *g*—i.e., for *ν* close to an eigenvalue of ***J*** —the recurrence plays a large role in shaping the dynamics.

Under this approximation, the plastic matrix converges to a static matrix during the stimulation period. Following stimulation for a time much longer than the synaptic timescale *p*, and given synapses slower than the inputs *p* ≫ 1*/f*, the plastic matrix is

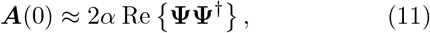

where *α* = *kI*^2^*/*4. Note the similarity of this expression and Eq. 4, the difference being that, in this case, **Ψ** is not an exact eigenvector of ***J*** . Nevertheless, this expression incorporates information about both the recurrent connectivity, ***J***, and the input structure both through **Ψ** and because the value of *ν* depends on the input’s amplitude and frequency (Methods).

We calculate the outlier eigenvalues of the matrix ***J*** + ***F*** (***J*** ) for a class of ***F*** (***J*** ) that includes the approximate ***A***(0) given by Eq. 11. Using a diagrammatic method, we show that if |*ν*| *> g*, then the outlier eigenvalues are (SM Sec. 2.3)

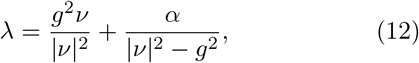

and its complex conjugate, unless |*λ*| ≤ *g*, in which case no outliers exist. Note that while the steps leading to the formula for ***A***(0) in Eq. 11 are approximate, given such a matrix, the above equation for the outlier eigenvalue pair is exact in the large-*N* limit. Eq. 12 is similar to Eq. 6 with *ν* playing the role of *η*; both have an *α*-dependent term contributing to the real part of the eigenvalue.

In the SM, we derive an additional condition describing the transition between the small oscillation and persistent oscillation regimes (Methods Eq. 19). This condition implies |*ν*| *> g* whenever oscillations persist, justifying our application of the above formula. We confirmed that Eq. 12 approximates the outliers observed in the simulations of Sec. II D (SM Fig. 8A) and that the transition boundary approximates what we observed in Fig. 2 (SM Fig. 9B). We emphasize that although this approximation captures the qualitative features of this model, it does not yield quantitatively exact results, particularly near the boundary between the small and persistent oscillation regimes (SM. Fig. 2).

We now trace the phenomenology of persistent oscillations for random-phase input back to our formula for the eigenvalue outliers. First, we note that, when *I* ≫ *g, k*— i.e., when the inputs dominate the recurrence—we find that |*λ*| *< g* (Methods), so that no outliers exist. Conversely, when *I* is excessively small, we find that our calculation correctly predicts that the network will be in the small oscillation regime (Eq. 19). This accounts for the observed amplitude-dependence of persistent oscillations: the amplitude must be weak enough to allow for dynamics to reflect contributions of the recurrence, but strong enough to avoid the small oscillations regime (Figs. 2Aii and 3).

Turning to the input frequency, we find that outlier eigenvalues only form within a preferred input frequency band. Within this band, the imaginary component of this eigenvalue outlier generally increases with the input frequency, which explains why faster inputs generally lead to faster persistent oscillations (Fig. 3C, SM Fig. 9A). These results emphasize that in this setting, the network learns to generate persistent oscillations that reflect the structure of the inputs, in contrast to the targeted inputs of the previous section.

Although the vector **Ψ** is not an eigenvector of ***J*** for |*ν*| *> g*, it overlaps with a small number of ***J*** eigenvectors (SM Sec. 3.3). To show this, we calculate the normalized overlap:

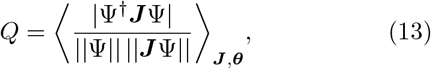

which is 1 when **Ψ** is an eigenvector of ***J*** and is 0 for a random **Ψ** which is uncorrelated with ***J*** . We show that *Q* → 1^−^ in the limit |*ν*| → *g*^+^, so that the vector **Ψ** becomes an eigenvector of ***J*** (SM Sec. 3.3). For intermediate values of *Q*, **Ψ** is partially aligned with an eigenvector of ***J*** ; such intermediate values of *Q* are found in the parameter regime in which oscillations persist (SM Fig. 9C). Numerically, we validate this by measuring the alignment between the leading eigenvectors of ***A*** and each eigenvector of ***J*** (Methods). As shown in Fig. 4A-Biv, we find that this projection localizes around a few complex eigenvalues in the persistent oscillation regime. Thus, the plastic matrix and neuronal activity aligns itself with eigenvectors of the static background, ***J***, in the persistent oscillation regime.

This is similar in spirit to what we found for targeted inputs with the key difference that the eigenvalue-eigenvector pair to which the network aligns itself is determined by the frequency and amplitude of the input. In a sense, the stimulus causes the network to express a pattern of activity, already embedded in the background connectivity, possessing dynamical properties that match those of the input. That is, the network anchors the input to pre-existing modes of the connectivity.

## III. DISCUSSION

We have shown that an ongoing Hebbian plasticity mechanism can learn to generate persistent oscillatory patterns of activity in response to both random and structured input. Classical models of working memory invoke Hebbian plasticity to explain persistent *static* activity that bridges temporal delays in working memory tasks [11–13, 30–32]. In these models, it is assumed that neurons with similar tuning profiles have excitatory recurrent connections between them, and this is justified by an appeal to Hebbian plasticity. Subsequent stimulus presentation leads to reverberating excitation between similarly tuned neurons, stabilizing the evoked activity patterns [31, 33]. Our work differs from these models in two important ways. First, the patterns of persistent activity in our model can be learned “on the fly,” in response to arbitrary input, rather than being burned into a prespecified connectivity scheme. Second, our model produces oscillatory rather than static persistent activity patterns. Many experimental studies report complex dynamical activity patterns following stimulus cessation during working memory tasks that do not align neatly with the picture of sustained firing [34–38]. Our results suggest that such complex persistent patterns could be generated by Hebbian plasticity processes that underlie canonical working memory models.

The vast majority of theoretical models of learning involve strong plasticity introducing entirely new dynamical patterns into a neural network [6–9]. In our model, plasticity affects ongoing neuronal dynamics by manipulating a dynamical reservoir provided by the static backbone connectivity. Previous work suggests that this could be the operating regime of many biological networks. For example, studies using brain-computer interfaces have found that neural circuits in motor cortex can only learn patterns of activity that lie within a constrained dynamical repertoire [39, 40], and studies in the hippocampus and sensory cortex have found that correlations in the activity patterns between neurons remain remarkably stable, even after periods of learning or across different task conditions [41–45]. These results suggest that plasticity and learning occur by adjusting pre-existing connectivity structures, rather than creating entirely new dynamical motifs from scratch. Our results emphasize that even microscopic changes in synaptic weights can have dramatic effects on neuronal dynamics and that interactions between pre-existing structure and ongoing plasticity can lead to novel forms of neural computation.

In our model, plasticity exerts a significant impact on network dynamics even though the size of the weight updates is small. Indeed, the elements of the plastic matrix *A*_*ij*_ scale as 1*/N*, while the individual baseline connections *J*_*ij*_ scale as 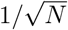. These small plastic updates can still influence neuronal dynamics when activity aligns with the low-rank structure of the plastic matrix [25]. For example, low-rank matrices with elements scaling as 1*/N* can produce order-one eigenvalue outliers when added to ***J*** (SM Sec. 2). In general, across different learning rules, there should be a relationship between the rank and the scale of plasticity-induced connectivity, an interesting topic for future study.

How might the persistent oscillations we describe relate to studies of oscillatory brain activity? Previous studies have reported persistent oscillations in neural activity following exposure to a periodic stimulus [46–50], and other studies have found that single neurons can phase lock to oscillations in the local field potential during working memory tasks [51–53]. These findings have sometimes been interpreted as evidence that the circuits being studied have an intrinsic tendency towards oscillation [46, 49]. Our results suggest that the situation may be more complex. Our networks have no a-priori oscillatory structure embedded in them, and in the random-phase case, no relationship to the stimulus. Yet, ongoing plasticity leads to persistent oscillations within preferred input frequency bands (Fig. 3C). A clear signature of such plasticity-induced oscillations is a decaying frequency over time (SM Fig. 5). This pattern cannot be explained by simple underdamped oscillator models. Observing such a slowing down in biological neural networks would suggest that plasticity induces persistent oscillations.

In the spirit of classical models of associative memory, we used targeted inputs to generate structured persistent oscillations [6]. More precisely, we aligned the phases of the oscillating inputs to a specific eigenvector of the background connectivity matrix, ***J*** . This process is similar to classical Hopfield models of associative memory in which a set of random patterns are embedded into a connectivity matrix and then evoked using pattern-aligned stimuli. In this way, the static backbone ***J*** encodes a “dynamical repertoire” of patterns that can be evoked through stimulation. We present evidence in SM Fig. 6 that this form of memory is extensive in the number of units.

The key difference between our model and Hopfield networks is that the synapses of our model are not held fixed. It is this feature that allows us to generate oscillatory, rather than static, memory patterns. Moreover, unlike Hebbian models for sequence storage [8, 9], our networks use *ongoing* plasticity to generate memory-related activity patterns, rather than discrete learning epochs. That is, we do not require learning episodes in which a pattern is written into the network prior to stimulation. Ongoing plasticity allows the network to generate persistent oscillations in response to inputs that the network has never seen before by anchoring the inputs to preexisting modes of the static connectivity (Sec. 1.6).

The Hebbian mechanism for persistent oscillations we studied may provide various functional advantages. Previous experimental work suggests that neuromodulators and other factors may gate or modify the strength of ongoing plasticity processes [54]. A previous study [25] examined static memory in the same network model we analyzed here. In a particular parameter regime in which the full network dynamics are chaotic, if synapses are frozen by halting plasticity, the network remains at a fixed point very close to the state it was in at the time of the freezing. In other words, any static state of the network can be held or remembered indefinitely by halting synaptic dynamics. Here we extend this form of memory across a richer dynamic range. Driving a similar network evokes a range of oscillations that persist indefinitely in the absence of input if synaptic plasticity is halted, freezing the synapses (Fig. 4E). This is essentially a form of input-driven reservoir computing in which a particular mode of the rich dynamics inherent in random network connectivity can be extracted by an input and then remembered indefinitely through the gating of plasticity. Such a computation could be useful for the maintenance of a dynamically extended memory or the generation of a rhythmic motor command. As another example of modulating plasticity, in SM Fig. 4, we altered *k* to induce persistent oscillations in networks that otherwise lie far outside the parameter regime in which this would otherwise occur.

Drawing from previous models of working memory [12], it would be interesting to design classes of ***J*** matrices from which ongoing plasticity can pull out more complex dynamical motifs through targeted stimulation. It may be that by choosing ***J*** appropriately, rather than randomly, one can design a network in which plasticity “pulls out” the appropriate patterns when cued with suitable transient stimulation. We leave this possibility to future work.

Overall, our work demonstrates that understanding memory-related neural activity may require modeling synaptic and neuronal dynamics together. Separating synaptic from neuronal dynamics, while convenient, can obscure basic functions of a circuit.

## IV. METHODS

### A. Simulation of plastic networks

We simulated the full neuronal-synaptic dynamics in Eqs. 1–2 using a fifth order Runge-Kutta (RK) solver with an adaptive step size. Specifically, we used the GPU-accelerated implementation of Tsitouras’ 5/4 method in Diffrax [55]. We set the absolute and relative tolerance of the solver to 10^−6^. In SM Fig. 3, we show that over a small subset of the parameters examined in the above figures, the lifetime and frequencies one obtains are left unchanged after increasing this precision by two orders of magnitude.

Unless stated otherwise, networks were run for 700 time units with the input lasting for the first 200 time units and with the parameters *p* = 25, *g* = 1.3, *f* = 0.1. In all analyses considered in the main text, the inputs *h*_*i*_ were present from the start point of the simulation. As shown in SM Fig. 2, this was not an essential choice. So long as an initial strong stimulation is given to the network to break it out of the small oscillation regime (Fig. 2A(ii)), the exact choice of initialization appeared to have little to no effect. Moreover, we show in SM Fig. 2 the lifetime of the persistent oscillations is unaffected by the amount of time the network is stimulated provided that the stimulation time is sufficiently long, a factor which scales with *p*.

### B. Chaos to entrainment transition

To calculate the entrainment to chaos transition line depicted in Fig. 2B, we used a bisection approach. For each value of *k*, we calculated the critical *I* above which more than half of a batch of 9 networks were fully entrained. We simulated networks of size *N* = 500 for 5,000 time units,with hyperparameters *g* = 1.3, *f* = 0.1, and *p* = 25. Networks were considered fully entrained when the average autocovariance function of the preactivations, Δ(*t* − *t*^*′*^) = ⟨*x*(*t*)*x*(*t*^*′*^)⟩, had a sinusoidal shape with no signs of decay between peaks. Numerically, we required that each peak of the autocovariance was within a distance of *ϵ* = 10^−3^ from all others. After obtaining critical *I* values for each *k*, we smoothed the resulting transition line using a cubic spline.

### C. Lifetime and frequency of persistent oscillations

We used the power spectrum of the network activity to measure the lifetime of persistent oscillations. Specifically, we calculated the average power spectrum of the preactivations over windows of width of 40 time units. A network was defined as oscillating at a certain time if the power spectrum of the preactivations within that window had a significant peak, defined as a point in the power spectrum that was 10^−1^ greater than adjacent points. We defined the lifetime of persistent oscillation as the time it took for the power spectrum to stop having any significant peaks. We show an example of this in SM Fig. 3.

We used a similar approach to measure the frequency of the persistent oscillations. Specifically, we calculated the power spectrum of the preactivations over a slightly smaller window of size 37.5 time units. The frequency of the ringing was then defined as the slowest frequency that had a significant peak in the power spectrum, where a significant peak was defined as above. If no such peak was detected, we simply set the ringing frequency to 0 in Figs. 3, 4, 5.

### D. Lifetime and frequency as a function of outliers

We obtained the results shown in Fig. 4C-D by considering a small 3 dimensional grid of *I, k*, and *f* values. For each combination of parameter values, we simulated a batch of 50 networks, each of size *N* = 500, and calculated the eigenvalues of ***J*** + ***A*** at the time the input was halted, as well as the lifetime and frequency of the resulting persistent oscillation. To obtain the results in Fig. 4, we first formed a 2 dimensional grid with the minimal and maximal *x* and *y* set equal to the minimum and maximum of the real and imaginary parts of the observed outlier eigenvalues, respectively. We then used Nadaraya-Watson smoothing with a Gaussian kernel to interpolate the lifetime and frequency of the persistent oscillations as a function of the outlier eigenvalue. Finally, because the outlier eigenvalue distribution was highly non-convex, we masked out points on the grid which were further than a distance of 0.1 from any observed eigenvalue. The results shown in Fig. 4 were generated by binning the resulting expected lifetime and frequencies.

### E. Eigenvalue outlier and transition formulae

In this section, we present the full formulae for the analytical results described in Sec. II F. See the SM for the full derivations. Recall that when an outlier eigenvalue pair forms in response to random-phase inputs, its value is given by

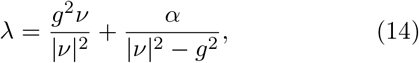

together with its conjugate. In this expression, the values of *α* and *ν* are given by:

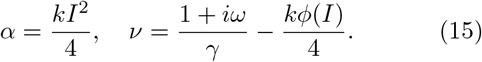

The value of *γ* is:

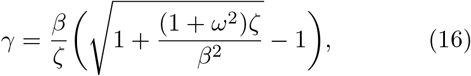

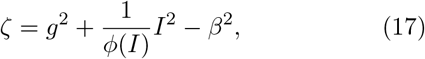

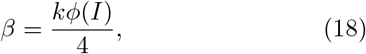

and *ω* = 2*πf* . We plot the behavior of *ν* as a function of the input parameters, *f* and *I* in SM Fig. 9A. Here, we make two observations based on the form of these equations. First, it is clear that the imaginary part of the outlier is determined by the term *g*^2^*ν/* |*ν*| ^2^. From the above expression, we can see that *ν* grows with *I*. This implies that the imaginary component of the eigenvalue will decay as inputs grow, explaining why we do not find persistent oscillations in response to strong inputs. Second, we note that the imaginary component of *ν* is given by *i*2*πf/γ*, and in many parameter regimes, Im(*ν*) grows with *f* (SM Fig. 9A). Thus, faster inputs generally lead to outliers with larger imaginary part. This explains why the frequency of persistent oscillations increases with the frequency of the inputs, within some admissible band (Fig. 3C).

In addition to this formula for the eigenvalue outlier *λ*, we approximate the transition from the small oscillation regime to the persistent oscillation regime. As described in more detail in the SM, the idea behind this calculation is to note that the constant component of the activity is typically large in the small oscillation regime and 0 in the persistent oscillation and input dominated regimes (Fig. 2A). We therefore identify the small oscillation regime with the parameter configurations that admit a non-zero time-independent component. Under our approximations and for ***J*** i.i.d. random with mean zero and variance *g*^2^*/N*, we find that this is equivalent to:

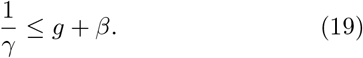

As shown in SM Fig. 9B, this approximates the transition between these two regimes well. Moreover, since 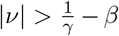, this implies that in the persistent oscillation regime, |*ν*| *> g*, which allows us to apply the results of SM Sec. 2.3 to estimate the eigenvalue outlier.

### F. Eigenvector overlaps

In Fig. 4(iv) we calculated the alignment between the plastic matrix, ***A***(*t*), and the eigenvectors of the random backbone, ***J***, at the time the input is turned off, *t* = 0. To do this, we first performed a spectral decomposition of ***A***(0), and we recorded the two eigenvectors with the largest eigenvalues, ***u, v*** ∈ ℝ^*N*^ . This was motivated by our analytical approximation in which ***A***(0) is exactly rank two. Next, we calculated the principal angle between the subspace spanned by ***u*** and ***v***, compared with the subspace spanned by the real and imaginary parts of each eigenvector of ***J*** . That is, given an eigenvector ***e*** = ***u***^*′*^ + *i****v***^*′*^ of ***J***, we calculated:

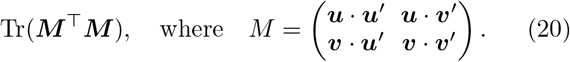

In Fig. 4(iv), we color each eigenvalue of ***J*** by the corresponding overlap value. This measure captures the alignment between the leading eigenvectors of ***A***(0) and each eigenvector of ***J*** .

### G. Targeted oscillations

For the targeting results we describe in Sec. II E, we drew 5000 different random backbone matrices *J* with *n* = 500. Then, for each random *J* matrix, we chose a random eigenvalue, formed oscillatory inputs following Eq. 7, and stimulated the networks for 200 time units, followed by a period without stimulation lasting 500 time units. For these analyses, we used *I* = 6, *k* = 3, *p* = 25, *f* = 0.1, and *g* = 1.3. After simulating the networks, we calculated the lifetime and frequency of the resulting persistent oscillation.

## CODE AVAILABILITY

Code and simulation data to reproduce all results is available at https://github.com/awakhloo/dynamic-memories [56].

## ACKNOWLEDGEMENTS

We thank members of the Aronov lab for feedback. This work was supported by the Gatsby Foundation (AW, DC, LFA), The Kavli Foundation (DC, LFA), and National Institutes of Health grants T32NS064929 (DC) and T32MH126036 (AW).

## Supplemental material

### 1 Additional simulations

**Supplementary Figure 1.**
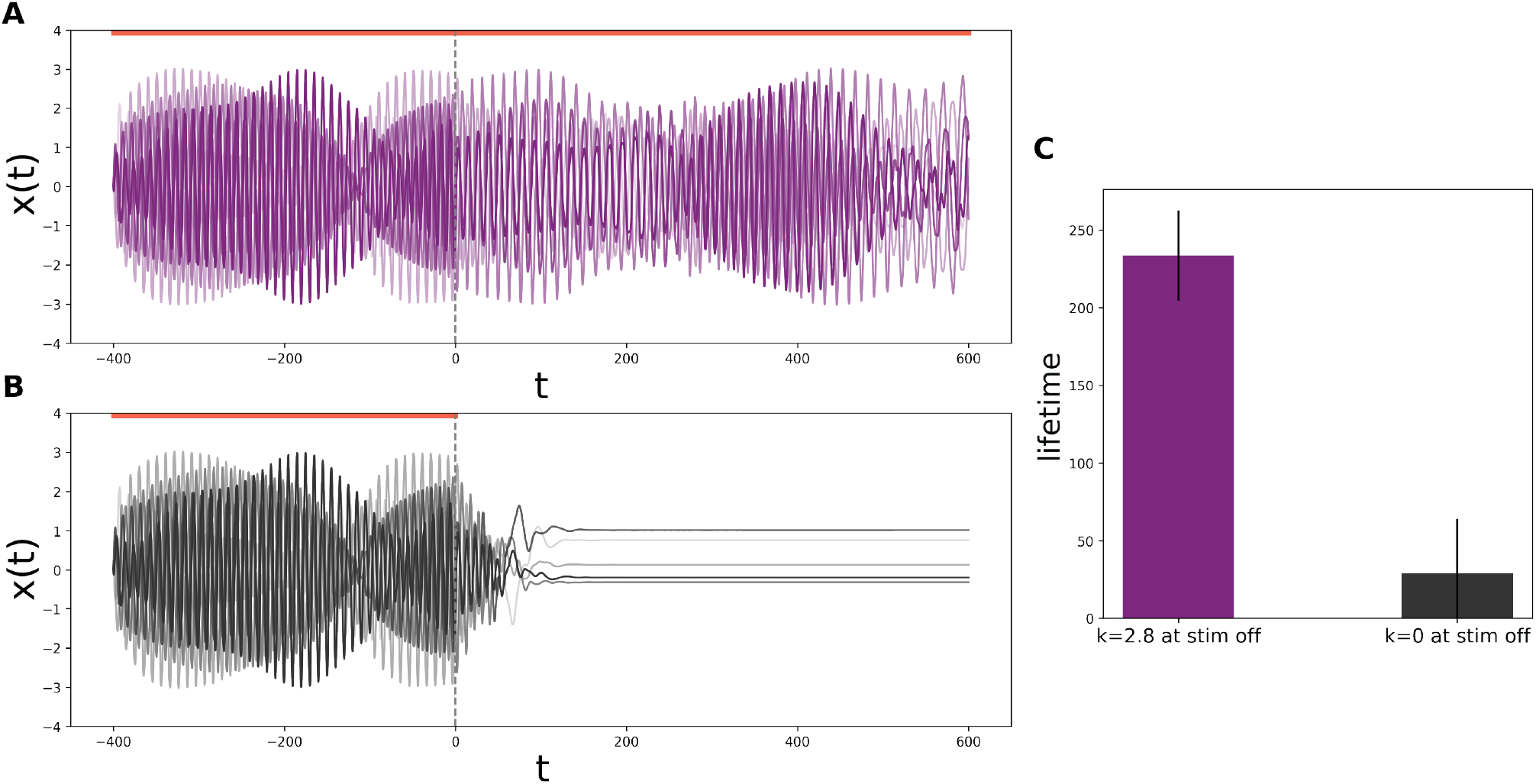
(A-C) Persistent oscillations are a self-renewing phenomenon. Here we show that persistent oscillations involve plasticity continuously reinforcing oscillatory neuronal activity after the stimulation is switched off. (A&B) Simulation of two networks with the same parameters (*k* = 2.8, *g* = 1.3, *p* = 25, *I* = 0.65, *f* = 0.1). In (A) the value of k is held fixed at *k* = 2.8 throughout, while in (B) it is set to zero when the stimulation is switched off (*t* = 0). Thus, in (B), the plastic weight matrix decays to zero exponentially at a time scale *p* = 25. In this case, oscillatory activity only lasts for a period which is *O*(*p*). (C) Here we rerun the paired simulations of (A) and (B) 50 times using different random backbone and input phases and compare the lifetime statistics between the two sets of networks. We plot the median lifetime in each condition, and error bars denote the standard error of the mean. Long-lasting persistent oscillations require ongoing synaptic plasticity after the input is switched off, demonstrating their self-renewing character.

**Supplementary Figure 2.**
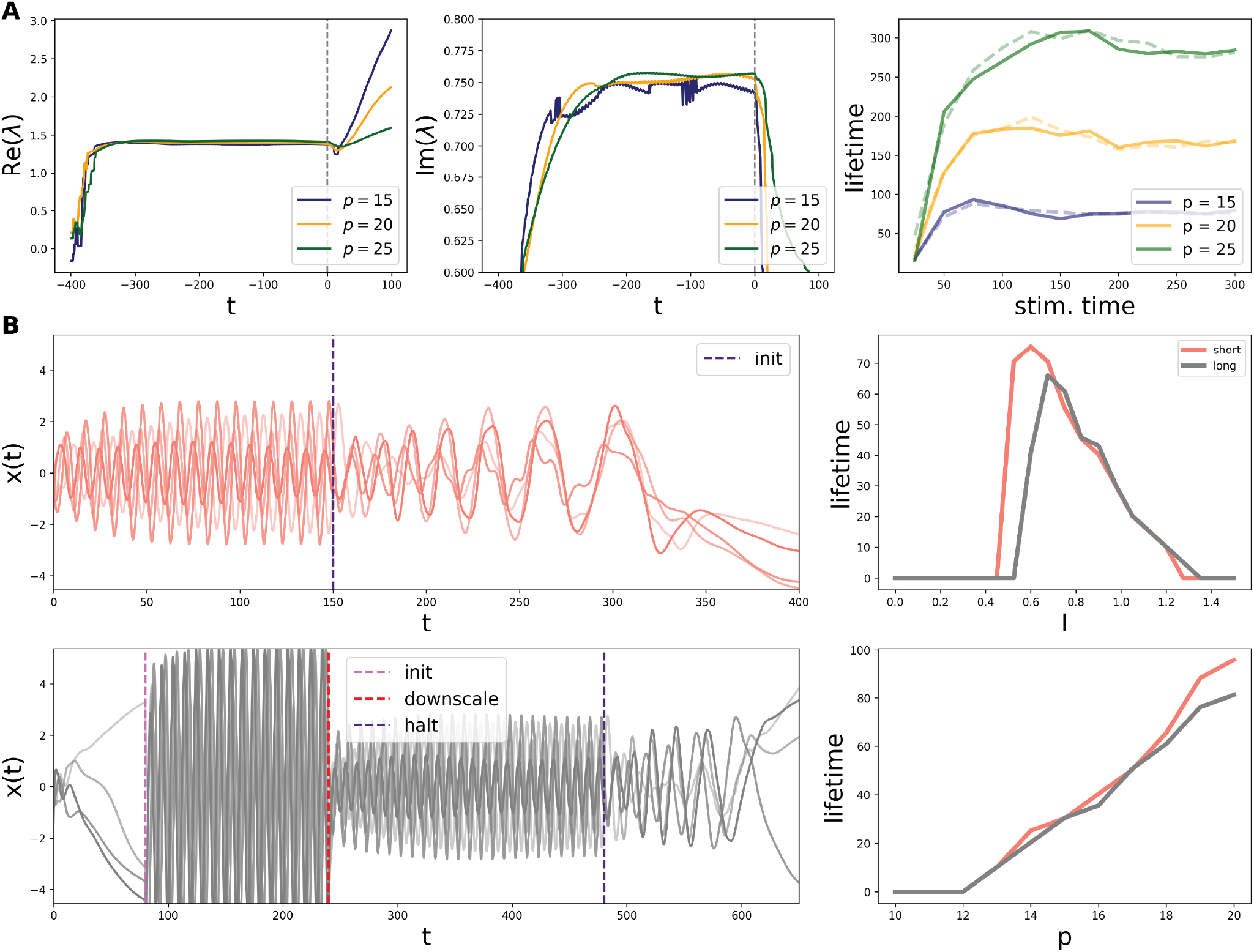
Effects of timing and initialization on persistent oscillations. (A) Dynamics of eigenvalue outliers during a stimulation period lasting 400 time units (left and middle), and the lifetime of persistent oscillations as a function of stimulation time, for various values of *p* (right) and with sizes N=1000 (dashed) and 5000 (solid). We use the parameters *k* = 3, *f* = 0.1, *I* = 0.75, and *g* = 1.3 for these analyses. (B) Initialization effects. Here we compare the lifetimes of two sets of networks: in the former (pink) networks, we stimulate the network from initialization for 150 time units. In the latter, (grey) networks, we allow the network to evolve without any input stimulation for 80 time units, followed by a strong stimulation period (init until downscale; *I* = 5), followed by a period of moderate stimulation (downscale until halt; *I* = 1). The initial large stimulation effectively breaks the network out of the small oscillation phase, allowing for persistent oscillations to occur. Note that unlike previous figures, time is not measured relative to the moment when stimulation is turned off–i.e., *t* = 0 at the start of the simulation. For both sets of networks, we initialize *x*_*i*_(0) according to a standard Gaussian, and we set *A*_*ij*_(0) = 0.

**Supplementary Figure 3.**
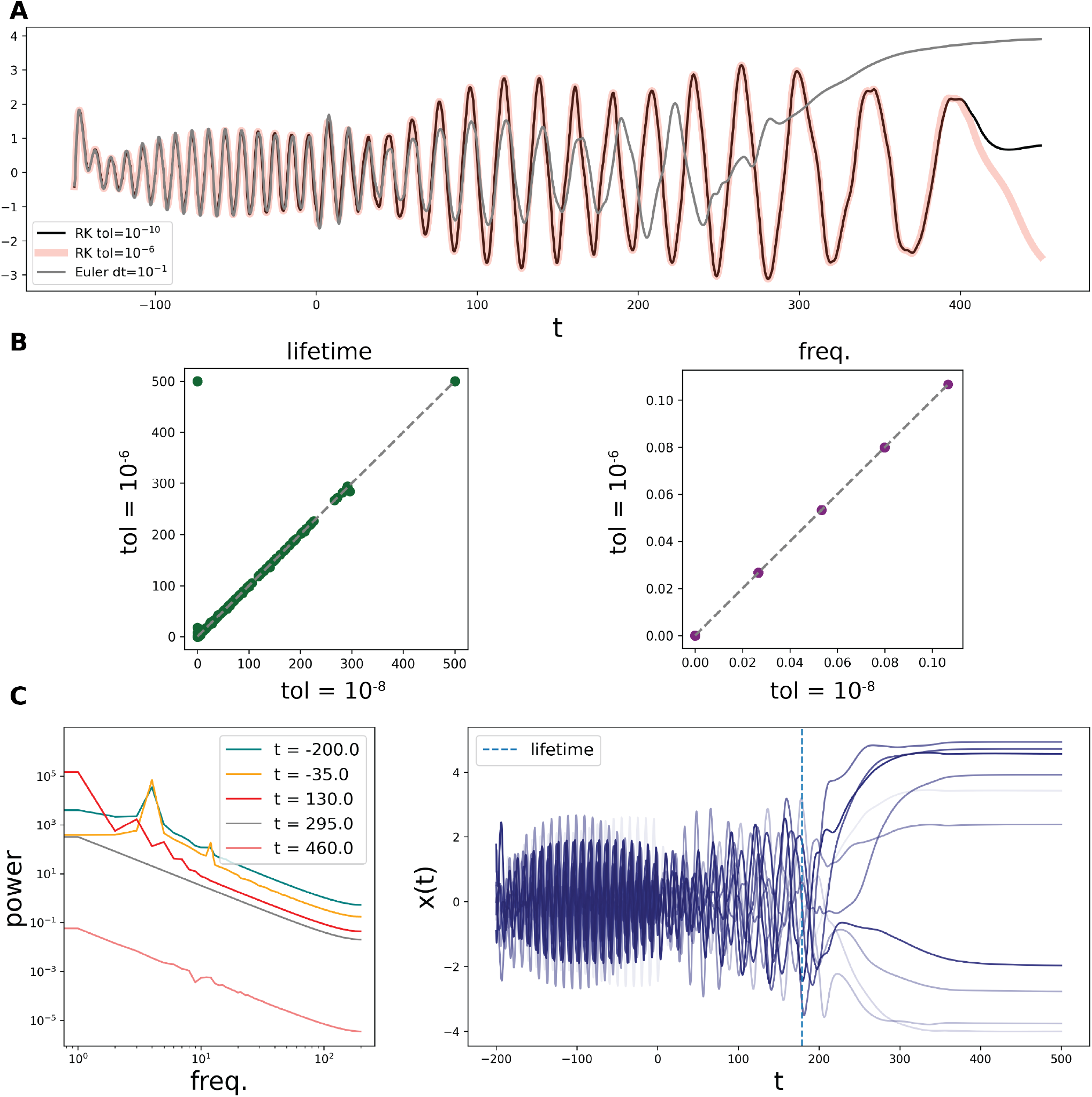
Validation of simulations. (A) Example network evaluated using a Runge-Kutta (RK) 5th order method [1] with high precision (black) as well as the precision used for the simulations in the main text (light pink), and an Euler method with *dt* = 0.1. The RK tol value corresponds to the maximum relative and absolute error. (B) Persistent oscillation lifetime and frequency statistics calculated using an RK method with the tolerance as in the simulations of the main text (y-axis), as well as a tolerance two orders of magnitude more precise (x-axis). We simulated 600 networks over a range of 60 parameter values using both sets of tolerances and found a close agreement between the resulting lifetimes and frequencies. (C) Example of our lifetime calculation method. We calculated the power spectrum of the network activity over sliding windows of size 37.5 time units and identified the first time at which the power spectrum did not exhibit a peak (Methods).

**Supplementary Figure 4.**
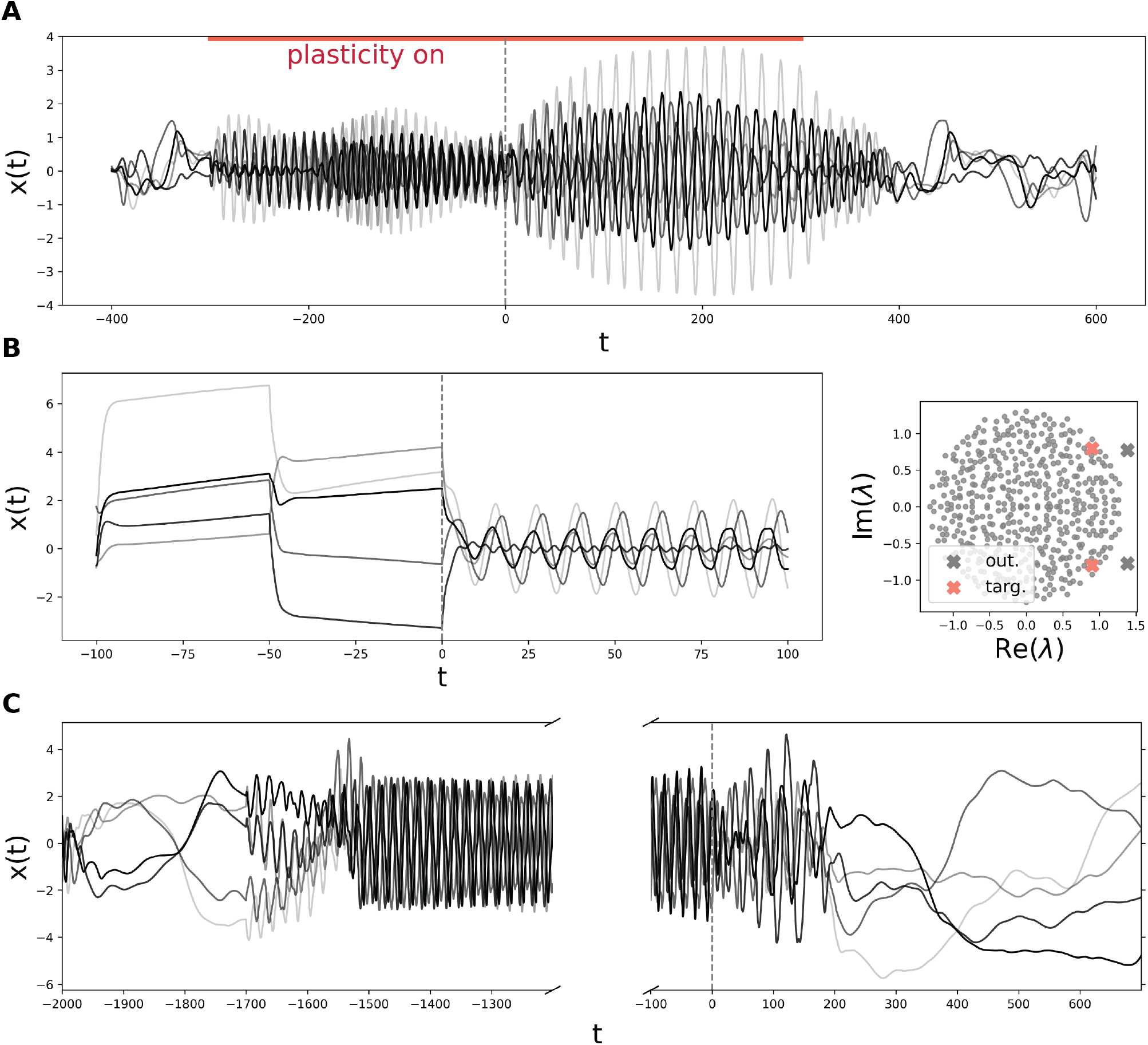
(A) Using a neuromodulatory signal to gate persistent oscillations. We initialize the network with *k* = 0, leading to spontaneous chaotic dynamics. We then stimulate the network with oscillatory inputs and set *k* = 3 (pink line). The input is switched off at the dashed line, and persistent oscillations occur. Finally, we set *k* = 0 once more at the end of the pink line, allowing the network to return to a chaotic state. This example demonstrates that a global neuromodulatory signal, reflected in the value of *k*, can be used to gate persistent activity in recurrent circuits. (B) Evoking persistent oscillations without oscillatory inputs. Here we consider a network with slow synapses (*p* = 100) and stimulate it with a single biphasic pulse aligned with an eigenvector, *ψ*_*i*_ = *u*_*i*_ + *iv*_*i*_. The input to each unit is set to *h*_*i*_(*t*) = *Iu*_*i*_ for 0 *< t <* 50 and is then set to the imaginary component, *h*_*i*_(*t*) = *Iv*_*i*_, for 50 *< t <* 100. Network parameters were set to *I* = 3, *k* = 2.5, *g* = 1.3, and we targeted the closest eigenvalue to *λ*_*targ*_ = 0.9 + 0.8*i*. (Left) Example units’ activity. (Right) Connectivity matrix, *J* +*A*(*t*) at the halt time, *t* = 0. The targeted eigenvalue pair (pink cross) is pulled out from the bulk through the stimulation period, leading to an outlier pair (grey cross). (C) Persistent oscillations in a parameter regime where the network activity is chaotic before and after stimulation. We set *N* = 1000, *k* = 1.9, *I* = 1, *g* = 1.7, *f* = 0.05, and *p* = 25, and we begin stimulating the network at *t* = −1700.

**Supplementary Figure 5.**
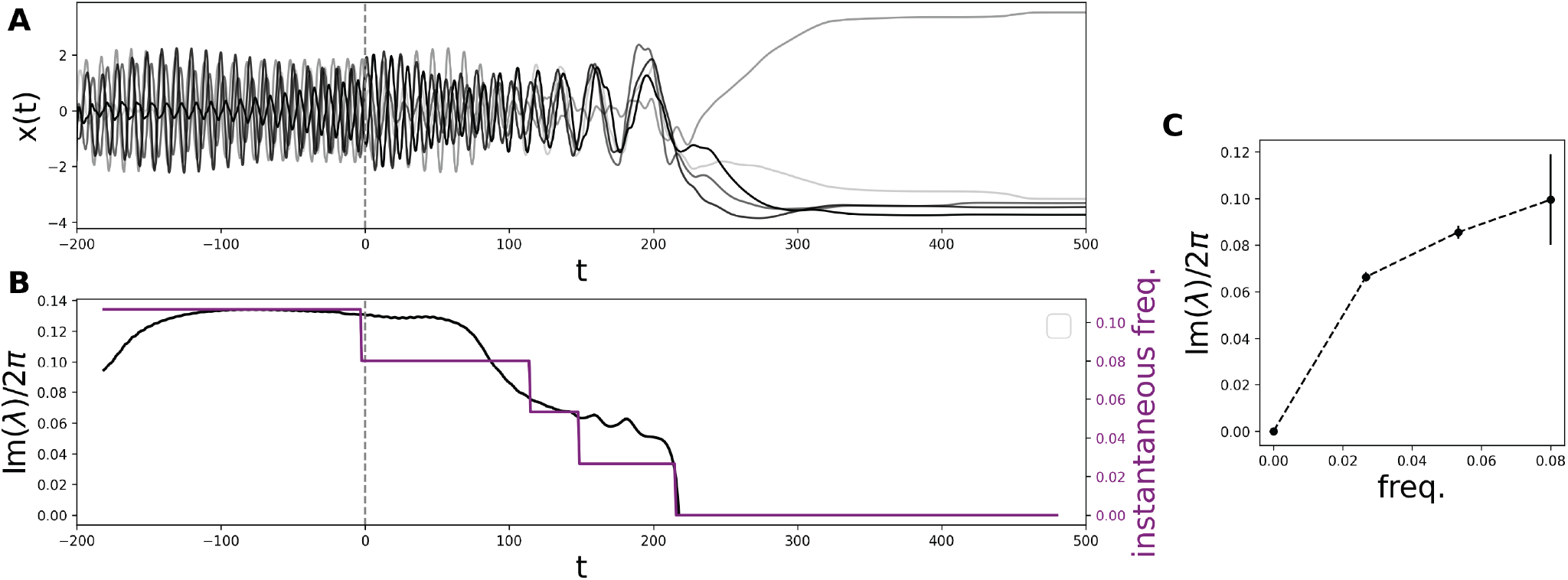
Instantaneous oscillation frequency tracks outlier eigenvalue. Here we show that the imaginary part of the eigenvalue outlier reflects the frequency of the network oscillations, following input cessation. (A) Example network trace. (B) Imaginary component of the eigenvalue outlier (black) and instantaneous network frequency (purple). The frequency is calculated by searching for a peak in the network activity’s power spectrum, calculated in windows of size *w* = 37.5. When no peak is detected we set the instantaneous frequency to zero. It is clear that in this example, the outlier dynamics closely track the instantaneous oscillation frequency. (C) Imaginary component of the outlier vs. instantaneous frequency. Here we simulated a set of networks over a small parameter grid and calculated the imaginary component of the outlier eigenvalue and the instantaneous frequency over the entire window following input cessation. We plot the mean ± SEM of the instantaneous frequency and the imaginary component of the outlier.

Here we target multiple eigenvectors in succession and show that even highly correlated eigenvectors evoke distinct patterns of activity. We first define two eigenvalue targets, *λ*_*targ*_, and, for each target, choose the two complex-conjugate eigenvalue pairs that are closest to these *λ*_*targ*_. We stimulate the network along each eigenvector for 200 time steps and then halt the input for an additional 200 time steps (Fig. 6A-B). For each eigenvector, we quantify the overlap of the activity with a given targeted eigenvector *ψ* as |⟨*ψ, x*(*t*)⟩|*/*||*x*(*t*)||, where the norm is taken over units, and we repeat this analysis across 40 different draws of *J* (Fig. 6C). We find that during the stimulation and persistent oscillation period, the activity is maximally aligned with the targeted eigenvector and has a non-trivial overlap with the eigenvector associated with the “neighboring” eigenvalue.

As shown in Refs. [2, 3], the eigenvectors corresponding to nearby eigenvalues are correlated with one another (Fig. 6D). This explains why there is a non-trivial overlap between the activity and the “neighboring” eigenvector. However, the fact that the network activity is maximally aligned with the targeted eigenvector shows that even highly correlated eigenvectors evoke distinct (persistent) oscillations. This suggests that the capacity of this form of memory grows with the number of distinct eigenvectors–i.e., it is extensive in *N* .

**Supplementary Figure 6.**
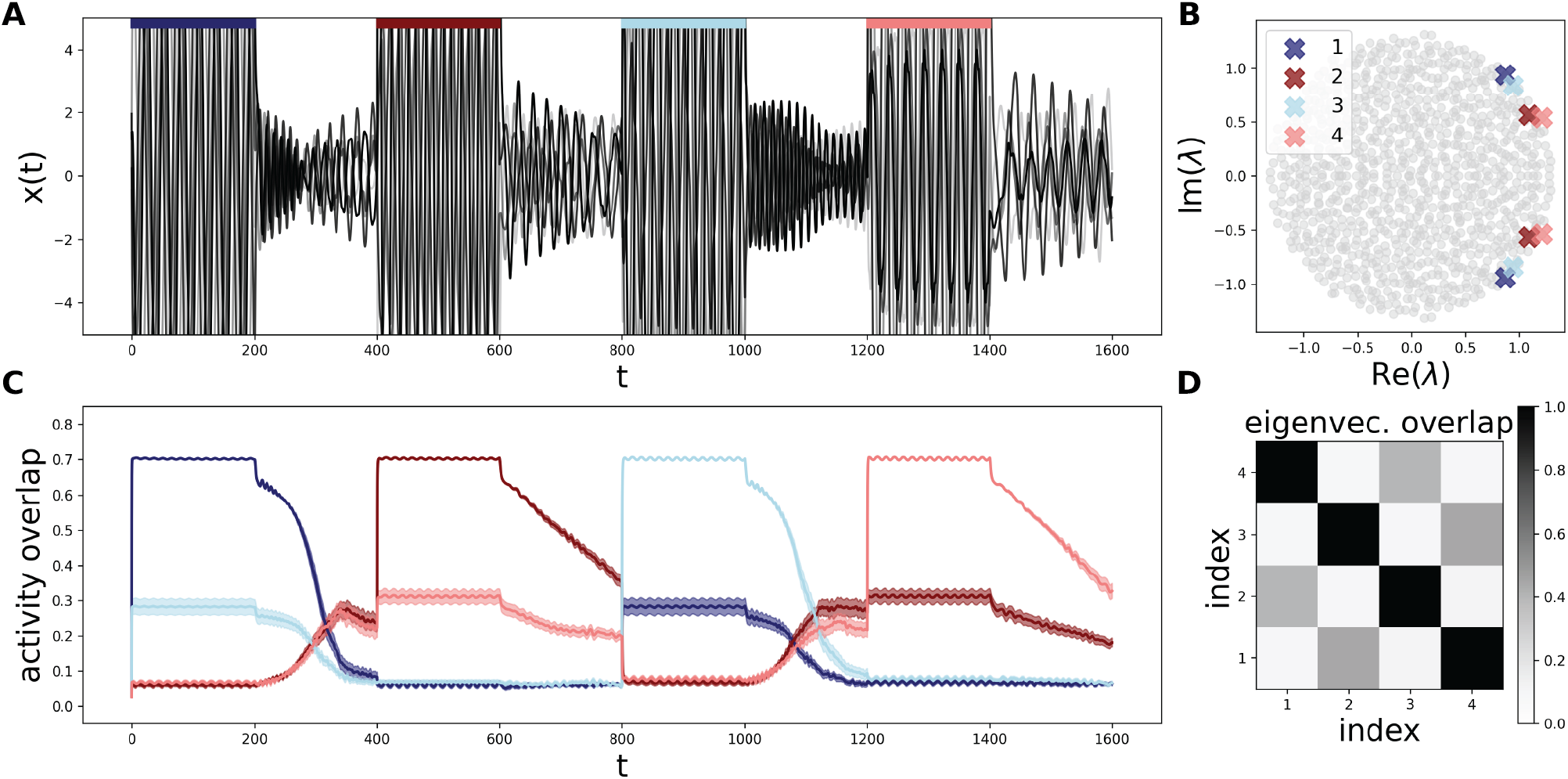
(A) Traces of 5 example units from the same network. Colored bars denote periods when we target the eigenvalue/eigenvector pair with the matching color. (B) Spectrum of *J* for the example network, with eigenvalue targets highlighted with crosses. Light/dark pairs correspond to neighboring eigenvalues–i.e., eigenvalues associated with the same *λ*_*targ*_. The legend denotes the order in which the corresponding eigenvectors are presented to the network. (C) Overlap of the preactivations with each eigenvector across 40 draws of *J*. During stimulation and early in the persistent oscillation period, the network activity is maximally aligned with the targeted eigenvector. Lines denote the mean and error bars the standard error across simulations. (D) Average overlap of the eigenvectors with one another. We use the absolute value of the inner product to quantify the overlap. Eigenvector indices are defined in terms of the order they are used to stimulate the network as in B. Across every simulation we set the *λ*_*targ*_ = 0.9+0.9*i*, 1.15+0.6*i* and choose the two closest conjugate eigenvalue pairs for each value of *λ*_*targ*_. Across every simulation, we set *N* = 1000, *g* = 1.3, *I* = 7, *f* = 0.035, *k* = 2.6 and *p* = 25.

### 2 Random matrix results

In the SM, we drop the main-text convention that vectors and matrices are bold. Additionally, we use the convention that matrix indices that do not appear on both sides of an equality are summed over.

In this section, we present the random matrix results described in the main text. The following results are exact in the large *N* limit and do not specifically apply to the neural network model considered in this paper. In the following section, Sec. 3, we employ a series of approximations which allow us to map the plasticity matrix *A*_*ij*_(*t*) onto one of the perturbations described below.

#### 2.1 Symmetric perturbation of a random asymmetric matrix

Consider a random *N* × *N*, asymmetric Gaussian matrix *J*_*ij*_ ∼ *N*(0, *g*^2^*/N* ). Let *H* be a symmetric matrix, *H* = *H*^⊤^, of rank *d* with real eigenvalues *λ*_1_, · · ·, *λ*_*d*_. Suppose further that *H* does not depend on *J* in the sense that ⟨*f* (*J*)*q*(*H*)⟩_*J*_ = ⟨*f* (*J*)⟩_*J*_ *q*(*H*), for arbitrary functions *f, q*. Then, in the limit *N* → ∞ with *d, λ*_*i*_ = *O*(1), any eigenvalue of *H* that lies outside the bulk of *J* will be an eigenvalue of the combined matrix, *J* + *H*. That is, the combined matrix *J* + *H* will have eigenvalue outliers at *λ*_*i*_ for any *λ*_*i*_ *> g*. This problem was studied rigorously in [4], but we give a heuristic derivation here, as we will use similar arguments in the following sections.

To show this, let *ξ* be an eigenvalue of *J* + *H*. Using the eigenvalue decomposition of *H*, one can write *H* = *O*Λ*O*^⊤^ = *UU*^⊤^ where Λ is diagonal, *O* is orthogonal, and *U* := *O*Λ^1*/*2^. By the matrix-determinant lemma we have:

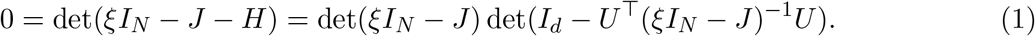

If *ξ* is an outlier eigenvalue, the first determinant is non-zero, so we focus on the second term. In the *N* → ∞ limit with *d* = *O*(1), this term is self averaging. In particular, to leading order in *N* :

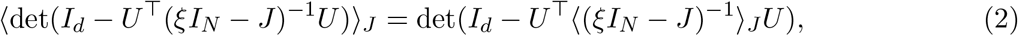

which follows after expanding the determinant about *I*_*d*_ − ⟨*U*^⊤^(*ξI*_*N*_ − *J*)^−1^*U* ⟩ and using the fact that *U*^⊤^(*ξI*_*N*_ − *J*)^−1^*U* concentrates about its mean in the large *N* limit with *d* = O(1). This can be shown by expanding the resolvent as a series in powers of *J* which is valid for |*ξ*| *> g* and averaging. In any case, using the formula for the resolvent, ⟨(*ξI*_*N*_ − *J*)^−1^⟩_*J*_ = 1*/ξ*, gives [5]:

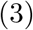

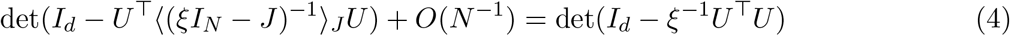

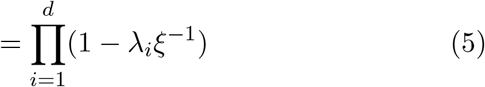

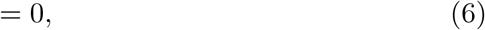

where *λ*_*i*_ are the (necessarily real) eigenvalues of *H*. It follows that there are outlier eigenvalues located at *λ*_*i*_ for |*λ*_*i*_| ≥ *g*.

#### 2.2 Eigenvector-aligned perturbation of a random matrix

We now consider eigenvector aligned perturbations of arbitrary, not necessarily random, matrices. Let *J*_*ij*_ be an arbitrary matrix, and let *ψ*_*i*_ be an eigenvector of *J*_*ij*_ with a corresponding eigenvalue *η* and unit norm, ∑_*i*_ |*ψ*_*i*_|^2^ = 1. Finally, let *α* be a real-valued scalar. We show that the combined matrix

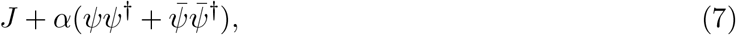

where the over bar is complex conjugation, has two eigenvalues located at

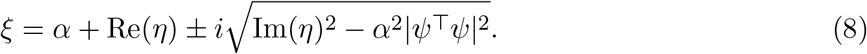

Taking *J*_*ij*_ to be random and Gaussian, we have the further simplification in the large *N* limit: *ξ* = *α* + *η*, together with its conjugate.

We show this using a similar line of reasoning as above:

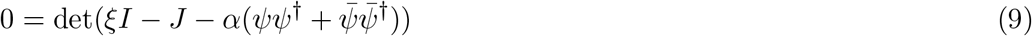

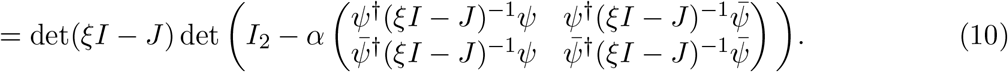

Once again, we focus on the second determinant, as we are interested in eigenvalues that form as a result of the perturbation, so we set

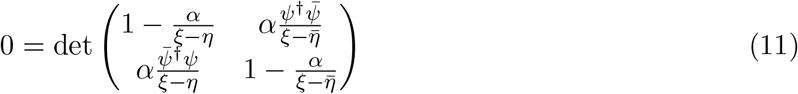

Solving for *ξ* gives the stated result

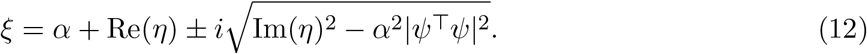

In the Gaussian setting, the rotation invariance of the distribution of *J*_*ij*_ suggests that in the large *N* limit, the real and imaginary parts of the eigenvector *ψ*_*i*_ are uncorrelated. Thus, the term |*ψ*^⊤^*ψ*|^2^ contributes a factor which is subleading in *N* . Expanding in the large N limit therefore gives the further simplification

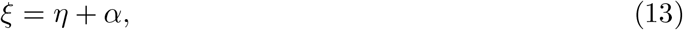

together with its conjugate.

#### 2.3 Resolvent-aligned perturbation of a random asymmetric matrix

Define the real symmetric matrix

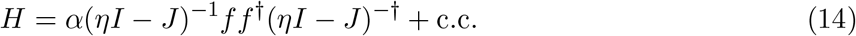

where *f* is an *N* -dimensional complex random vector with i.i.d. entries satisfying: ⟨*f*_*i*_⟩ = 0 and ⟨|*f*_*i*_|^2^⟩ = 1*/N* . Let *J*_*ij*_ ∼ *N*(0, *g*^2^*/N* ) be a random asymmetric Gaussian matrix. Finally, assume that |*η*| ≥ *g*. Under these conditions, in the large *N* limit, the combined matrix *J* + *H* has outlier eigenvalues located at

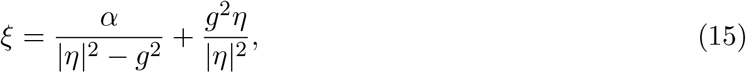

together with its conjugate, if the right hand side lies outside the bulk: |*ξ*| ≥ *g*. In Sec. 3.2, we show how the connectivity matrix can be mapped onto this setting when the network is stimulated with random inputs.

To derive this, we again start from the equation det(*ξI* − *J* − *H*) = 0. Following an analogous line of reasoning as above, one eventually obtains, after averaging over *f* :

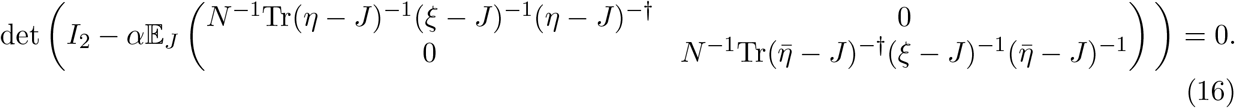

Using the fact that the resolvent is analytic for |*η*|, |*ξ*| ≥ *g*, we can expand the trace in the upper-left entry as

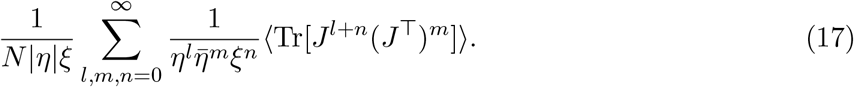

We now show that in the large *N* limit only the diagonal terms, *m* = *l* + *n* contribute, and that the contribution of the trace terms is *Ng*^2*a*^. Formally, we show:

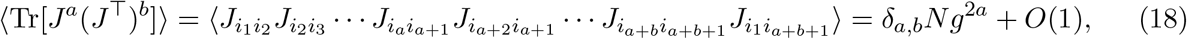

where we have again used the convention that matrix indices that do not appear on both sides of an equality are summed.

We derive the formula above by showing that at most a single Wick pairing for each *a* and *b* survives in the large *N* limit. This is done using a diagrammatic method: each matrix index is associated with a vertex, and shared indices in the trace above are connected by a line (SM Fig. 7A-B). Diagrams corresponding to individual Wick pairings are generated by connecting the left and right indices to one another–e.g. by connecting *i* to *j* and *j* to *k* in the term ⟨*J*_*ij*_*J*_*jk*_⟩. Indices that are shared by two *J* terms (i.e. *i*_2_, …, *i*_*a*_) are drawn on top, those shared by two *J*^⊤^ matrices at the bottom, and the indices connecting *J* to *J*^⊤^ (i.e. *i*_1_, *i*_*a*+1_) are drawn vertically (see SM Fig. 7B). Indices appearing in a closed loop are summed over.

We now consider how each diagram scales in the large *N* limit. In this notation, the number of matrices involved in the product, *a* + *b*, is given by the number of horizontal lines. Since each set of double lines corresponds to a tensor *g*^2^*N*^−1^*δ*_*ik*_*δ*_*jl*_, each loop corresponds to a sum over a unique index which brings a factor of *N* . Thus, each diagram scales as *N* ^#loops−#gaps*/*2^. Our claim therefore reduces to the statement that there is at most one non-vanishing diagram which exists only for *a* = *b* and that its value is *g*^2*a*^*N* .

The dominant diagrams are those in which the horizontal lines are paired off, as in SM Fig. 7B and C. To show this, it suffices to note that the number of loops is maximized by the planar diagrams, as any crossing of lines leads to the formation of a larger loop (see, e.g., [6, 7] for more thorough discussions of the role of planar diagrams in Hermitian and non-Hermitian rando matrix theory). In the expansion considered here, there is only one such pairing, corresponding to the diagrams shown in SM Fig. 7B and C. The fact that this pairing is only possible when *a* ≠ *b* shows that the *a* ≠ *b* terms are subleading, and shows that for *a* = *b* there is a single dominant diagram at each order which is equal to *g*^2*a*^*N* .

**Supplementary Figure 7.**
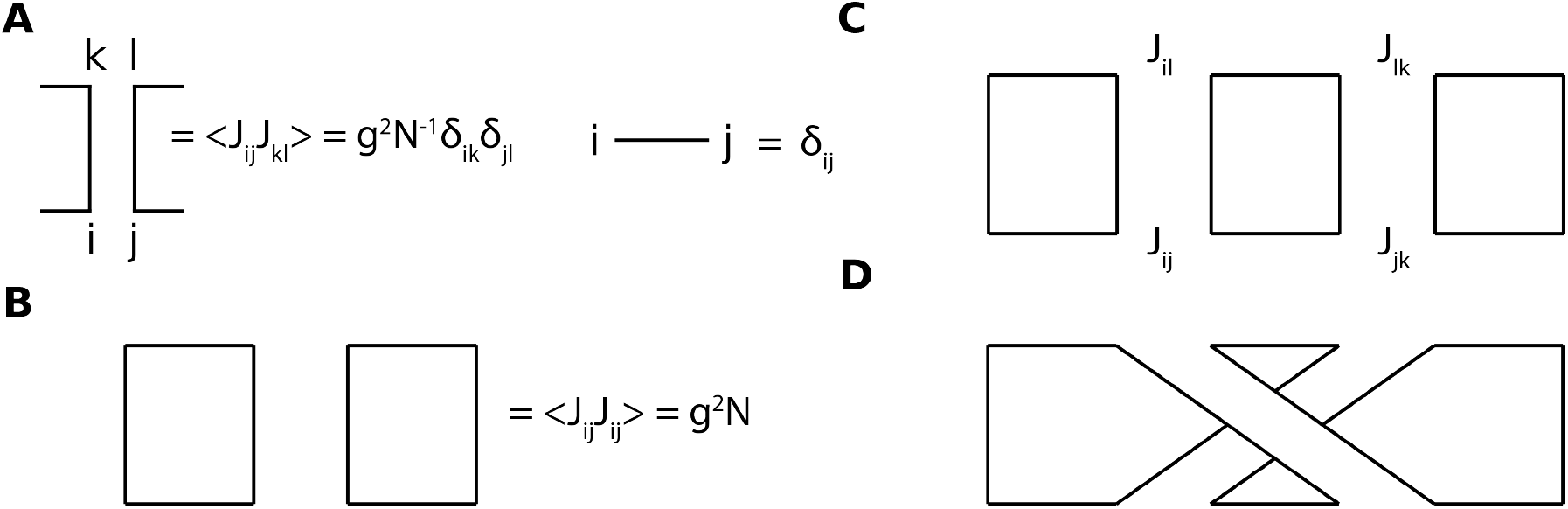
(A) Diagram rules. Double lines are used to denote specific Wick pairings. (B) Example diagram corresponding to the term Tr(*JJ*^⊤^) = ⟨*J*_*ij*_*J*_*ij*_⟩ . (C-D) Two example diagrams appearing in the expansion of Tr[*J*^2^(*J*^⊤^)^2^]. The diagram in D corresponds to the pairing ⟨*J*_*ij*_*J*_*lk*_⟩⟨*J*_*jk*_*J*_*il*_⟩ consisting of a single loop and is subleading, while the one in C corresponds to the term ⟨*J*_*ij*_*J*_*il*_⟩⟨*J*_*jk*_*J*_*lk*_⟩ and is an example of a dominant diagram which scales as *O*(*N* ).

We are therefore left with the sum:

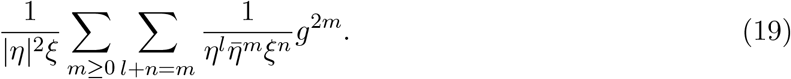

We now carry out the sum over *l, n* by recognizing the geometric series:

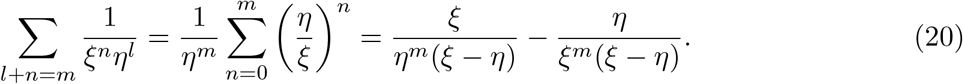

Carrying out the remaining geometric sums and simplifying then gives

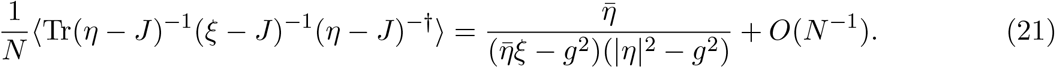

Inserting this into Eq. (16), we obtain the stated formula for the outlier eigenvalue:

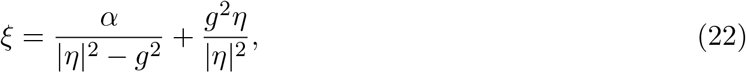

together with its complex conjugate.

### 3 Eigenvalue outlier formulae

In the following sections, we employ a number of approximations to obtain analytical expressions for the neuronal-synaptic dynamics considered in the main text. The result of these approximations is a functional form for the plastic matrix, *A*_*ij*_(*t*) following a sufficiently long stimulation period. With this functional form in hand, we can apply the results of the preceding section to obtain formulae for the eigenvalue outliers of the connectivity matrix.

#### 3.1 Targeted inputs

In this section we derive the formula for the eigenvalue outliers in the targeted inputs case. We start with this case, since it is simpler than the calculation for random inputs. The dynamical equations in the targeted setting are

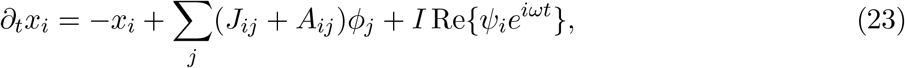

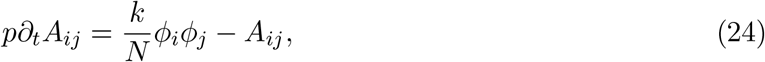

where we have used the shorthand *ϕ*_*j*_ := *ϕ*(*x*_*j*_), and the vector *ψ* is an eigenvector of *J*. We calculate the value of the eigenvalue outlier in the limit of strong inputs and slow synapses and inputs: *I, p* ≫ 1*/ω* ≫ 1, as well as a long stimulation time, *t* ≫ *p* ≫ 1. In this limit, the neuronal dynamics during the stimulation period are completely input driven. To leading order in *I* and *ω* we have

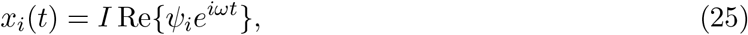

In this limit, the plastic matrix converges to a static value: *A*_*ij*_ = ⟨*ϕ*_*i*_*ϕ*_*j*_⟩, with brackets denoting time averaging, following a sufficiently long stimulation time, *t* ≫ *p* ≫ 1. Defining the phase of the input, *θ*_*i*_ := atan2(Im[*ψ*_*i*_], Re[*ψ*_*i*_]), we can calculate the entries of *A* as

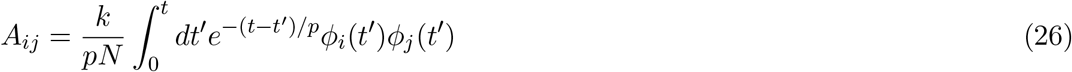

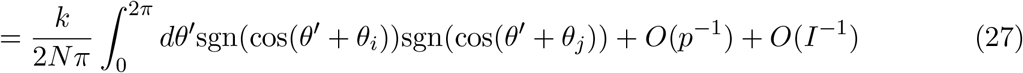

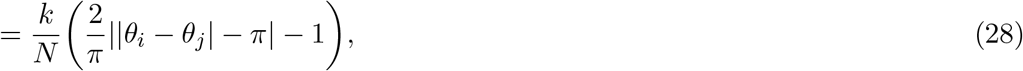

where sgn is the signum function, and we have used the fact that |*θ*_*i*_ −*θ*_*j*_| ≤ 2*π* in passing to the last line. The plastic matrix may therefore be written as a function of the input phases: *A*_*ij*_ = *A*(*θ*_*i*_−*θ*_*j*_). We now use this fact to calculate the spectral decomposition of *A*. The eigenvector equation is

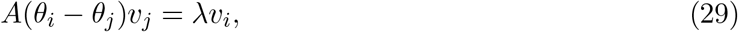

where summation is implied. Replacing sums with integrals in the *N* → ∞ limit, we have the eigenfunction equation

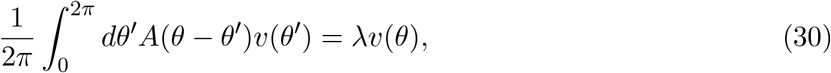

It follows that the eigenfunctions are sinuisoids. Moreover, since the above equation has a phase symmetry *v* ↦ *e*^*iφ*^*v*, the eigenfunctions will be pairs of cosines and sines up to an arbitrary phase shift. To leading order, the eigenvectors are then a discretized version of these eigenfunctions with unit norm:

**Supplementary Figure 8.**
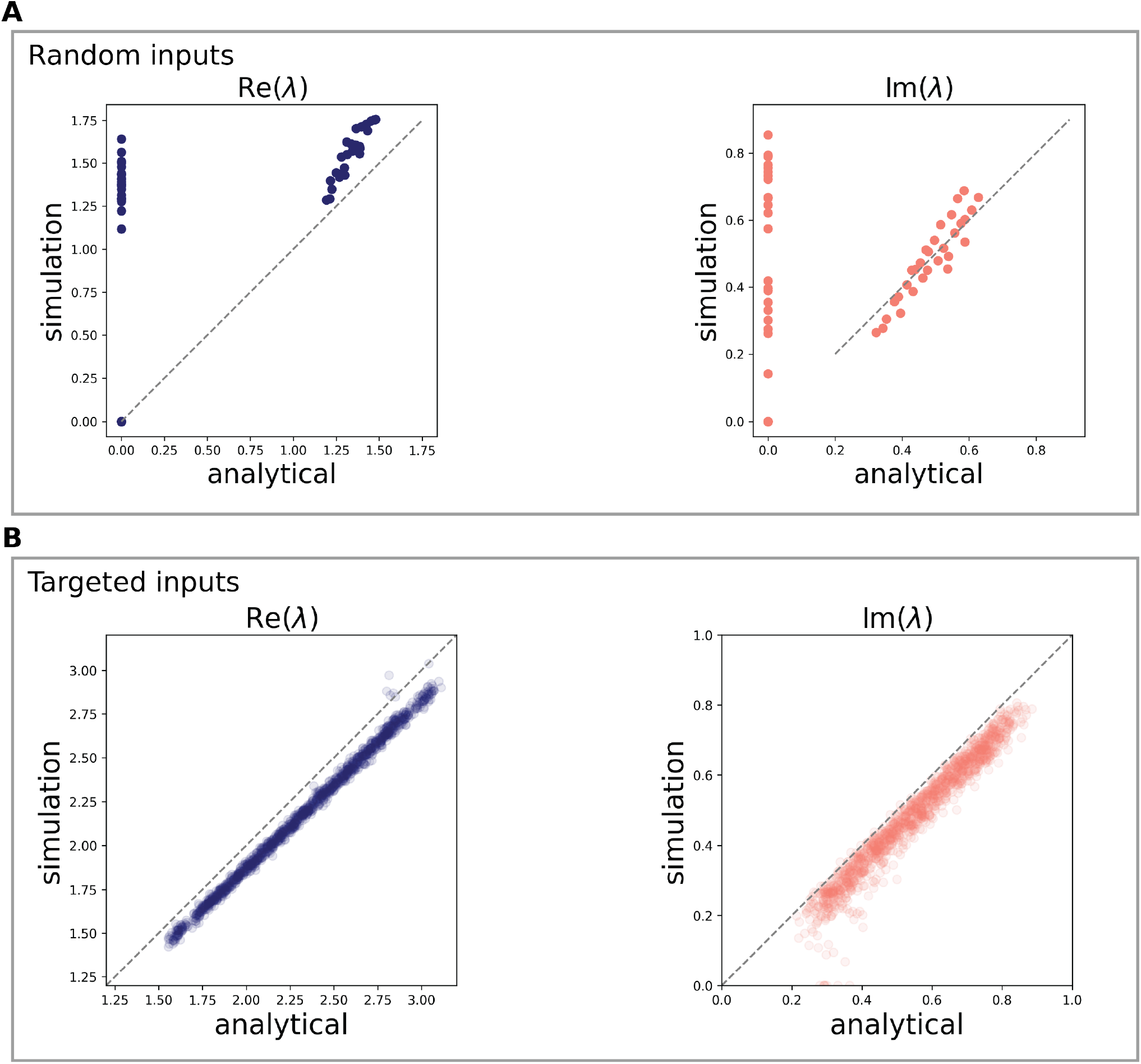
Match between analytical and empirical eigenvalue outliers. (A) Random phase case. Here, we compare the outlier eigenvalues obtained in the simulations for Fig. 3 with the predictions from our formula, Eq. (55). Each marker corresponds to an average over 50 simulations. Note that the accumulation of values on the y-axis correspond to parameter settings that produced eigenvalue outliers in spite of the fact that one of the analytical conditions for an outlier were not met (|*λ*| ≤ *g* or the small oscillation transition described in Eq. (46)). Our calculations approximate the outlier well, but is excessively aggressive in detecting the small oscillation transition. (B) Targeted phase case. Here, we sweep over a range of *k* ∈ [2, 5] and eigenvalue target (*η*) values, as well as a smaller set of *I* and *f* values, and plot the resulting match between simulation and our formula, Eq. (40).

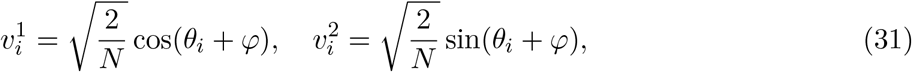

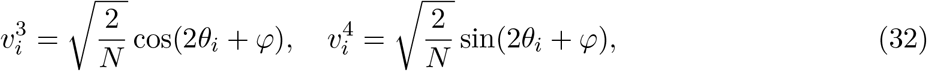

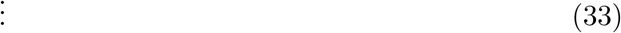

We can now calculate the eigenvalues to leading order in *N* as:

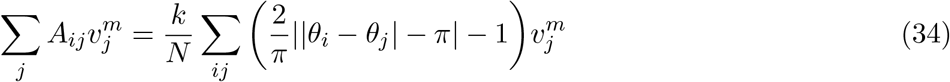

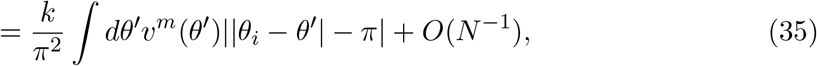

from which it follows that the eigenvalues come in pairs. The *m*th pair is given by

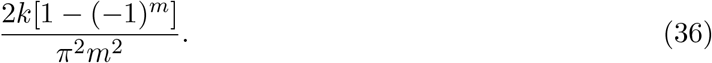

Given the fast decrease in the eigenvalues, we can form a rank two approximation of *A* by truncating after the first eigenvalue pair. That is, we set

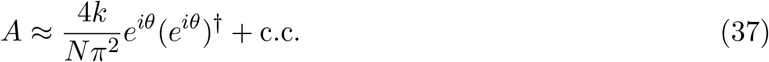

With this closed-form solution for *A* in terms of the inputs in hand, we can proceed to calculating the eigenvalue outlier.

We first decompose *J* as

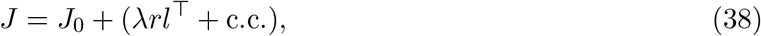

where again *r* is the eigenvector that we target and *l* is the corresponding left eigenvector. To make progress, we now approximate *J* with the matrix

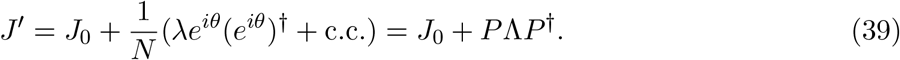

Essentially, we have replaced the eigenvector *r* and its conjugate with a pair of eigenvectors whose entries have absolute value 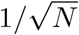. In this way, the matrix *A* in Eq. (37) becomes perfectly aligned with the eigenvector of *J* that has eigenvalue *λ*. This approximation incurs a small bias in the resulting eigenvalue equation (SM Fig. 8B), but it captures the trends very well. We can now use the result of Sec. 2.2 to calculate the outlier eigenvalues of the combined matrix, *J*^*′*^ + *A* with *α* = 4*k/π*^2^. This gives

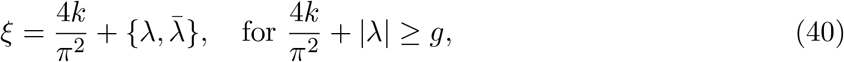

reproducing the equation given in the main text.

#### 3.2 Random inputs

We now consider the case of random inputs in the limit, *p* → ∞. As in the case of structured inputs, we begin by obtaining an approximation of *A*, which again converges to a static matrix in the large *p* limit.

We begin by assuming that, at any point in time during the stimulation period, the vector of post-activations *ϕ*_*i*_(*t*) is an eigenvector of *A*:

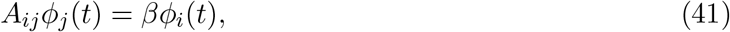

where *β* will be determined self-consistently. This assumption is motivated by the form of the Hebb rule: given a collection of *N* time-varying sinusoids with random phases and amplitudes *g*_*i*_(*t*), the matrix *C*_*ij*_ = ⟨*g*_*i*_*g*_*j*_⟩_*t*_ will satisfy *C*_*ij*_*g*_*j*_ = *βg*_*i*_ to leading order in *N*, with the exact value of *β* depending on the distribution of amplitudes of the *g*_*i*_. In the parameter regime in which persistent oscillations occur, the post-activations *ϕ*_*i*_ are approximately sinusoidal, motivating the above assumption.

The neuronal dynamics reduce to:

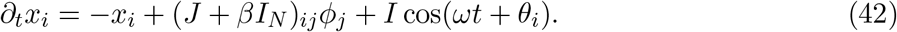

We now make the further simplifying assumption that the non-linearity acts only by distorting the scale of the pre-activations, without affecting their shape:

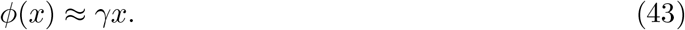

We expect this approximation to describe the network activity when most units are far away from saturating the non-linearity. Moreover, even when many of the units *do* saturate the non-linearity, this approximation can be motivated by the fact that the dynamics of the units are strongly input-driven in the persistent-oscillation regime–i.e., they are highly oscillatory. This suggests that there exists a *linear* recurrent neural network, driven by external oscillatory input, whose macroscopic statistics match those of the full, non-linear network. The goal of this calculation is therefore to find values of *β, γ* which yield such a linear network. Note that while in the previous section we appealed to a large input amplitude *I* ≫ 1 limit, this is not possible here, as persistent oscillations do not occur in the large *I* regime.

Under these assumptions the neuronal dynamics can be solved exactly,

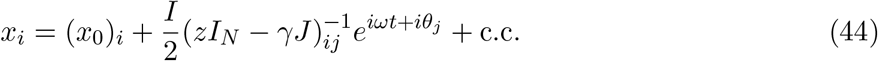

Where we have written the neural activity as the sum of a static component *x*_0_ and an oscillatory component. Furthermore, we defined *z* := 1 + *iω* − *βγ*.

We begin by considering the static component of the activity, which must satisfy by Eq. (42):

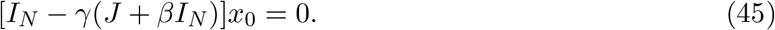

In the persistent oscillation regime the static component of the activity is 0, while in the small oscillation phase it is large. Under the linear approximation, the transition between these two states is governed by the existence of a time-independent component of the activity–i.e., the existence of a solution, *x*_0_, to the above equation. Given that the eigenvalues of *J* uniformly fill a disk of radius *g*, this equation will typically have a solution valid to 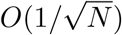 when

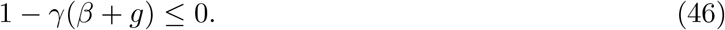

In what follows, we will assume that we are in the persistent oscillations regime, so that the inequality above does not hold, and thus, *x*_0_ = 0. We will thus enforce 1 − *γβ* − *γg >* 0 as a necessary condition for an outlier eigenvalue to form. This ensures that outlier eigenvalues are not predicted for parameter settings that lead to small oscillations (SM Fig. 9).

We proceed by fixing the value of *γ*. We do this by imposing that the mean square amplitude of the *ϕ*_*i*_ saturate at ±1 for large input amplitudes *I*. More precisely, let us define the amplitudes of the *x*_*i*_ as 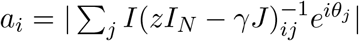 and their mean-square as *ā* = *N*^−1^ ∑_*i*_ |*a*_*i*_|^2^. The corresponding mean-square *ϕ*_*i*_ amplitude is given by *γ*^2^*ā*. We now set *γ*^2^*ā* = *ϕ*(*I*), so that this amplitude saturates at 1 for large *I*, matching the model phenomenology. This choice gives the condition:

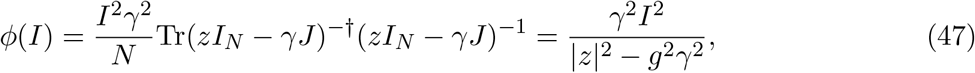

where we have assumed |*z/γ*| ≥ *g* and expanded the resolvent as a series in *J* to simplify the trace along the lines of the derivation in Sec. 2.3. Solving for *γ* gives

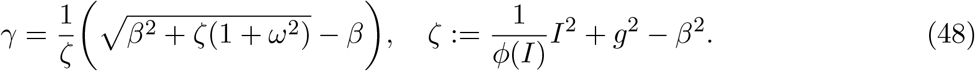

The eigenvalue *β* now follows from *ϕ*(*x*) ≈ *γx*. This allows us to write:

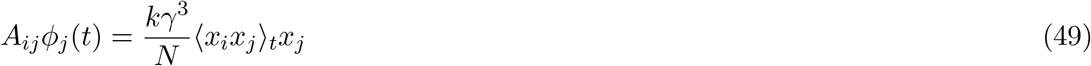

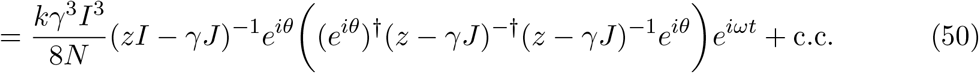

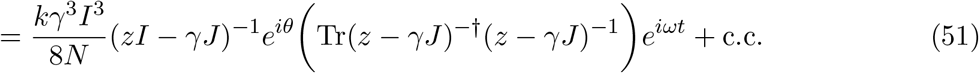

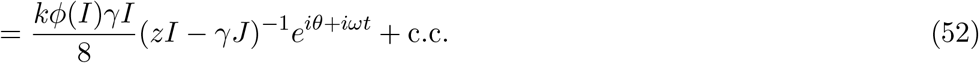

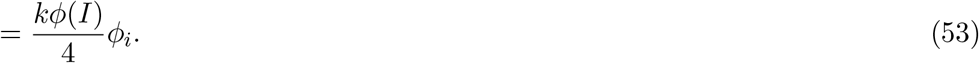

where we have used Eq. (47) to simplify the trace and have dropped terms subleading in *N* . It follows that *β* = *kϕ*(*I*)*/*4. Note that this line of reasoning is consistent with our starting assumption that the vector *ϕ*_*i*_ is an eigenvector of the plastic matrix *A*_*ij*_.

Taking *p* ≫ 1*/ω*, the plastic matrix then converges to the static matrix

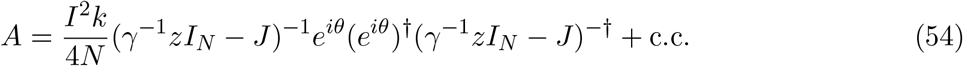

Using the result in Sec. 2.3 above then gives the eigenvalue outliers as a function of the model hyperparameters. Defining *α* := *I*^2^*k/*4 and *ν* := *z/γ*, we have that the outlier *ξ* is given by

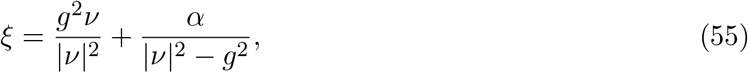

together with its conjugate, unless |*ξ*| ≤ *g* or Eq. (46) holds. Note that since |*ν*| *> γ*^−1^ − *β*, the latter condition implies that |*ν*| *> g* in the persistent oscillation regime, justifying our application of the results of Sec. 2.3.

We now show that this solution captures the key input sensitivity characteristics of persistent oscillations for random inputs (SM Fig. 9). With regards to the amplitude, in the strong input limit *I* ≫ *k, g*, we find *γ* ∼ 1*/I*. In this limit, it follows that |*ν*| → ∞, and thus, the imaginary part of the outlier, 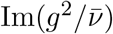, tends to zero. Conversely, in the weak input limit, *I* → 0^+^, we find *β* → 0 and *γ* → |1 + *iω*|*/g*. By Eq. (46), it follows that this leads to the small oscillation regime, as expected (SM Fig. 9). This captures the fact that neither excessively large, nor excessively small inputs can produce eigenvalue outliers, and thus, persistent oscillations (Main Fig. 3). With regards to the input frequency, this solution explains another key feature we observed in the model: within the admissible frequency band, faster inputs lead to faster persistent oscillations (Main Fig. 3). From this calculation, we find that faster inputs lead to larger imaginary components of *ν*, and thus larger imaginary components in the outlier *ξ*, when it exists (SM Fig. 9A). Taken together, these results show that our linear approximation captures the key behaviors of the model.

#### 3.3 Eigenvector alignment with random inputs

In Section 3.1, we showed that in the targeted setting, the plastic *A* matrix becomes aligned with the targeted eigenvector of *J*, and this leads to the formation of complex outliers. In this section, we show that a similar phenomenon occurs using random inputs: when persistent oscillations occur, the neuronal dynamics spontaneously align themselves with eigenvectors of *J*, and this leads to the formation of complex outliers via the plastic matrix, *A*. This alignment occurs in spite of the fact that the input phases are chosen randomly, without any knowledge of *J*.

To begin, let us write the approximate value of *A* in response to random inputs (Eq. 54) as

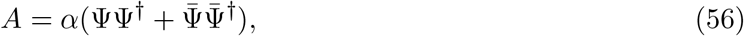

where 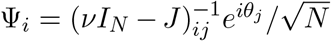. We show that as |*ν*| → *g*^+^, the vector Ψ becomes an eigenvector of *J*. Thus, in this regime, multiplying the random vector of phases 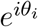, by the resolvent matrix yields an approximate eigenvector of *J*. In the random phase case, persistent oscillations occur in an intermediate regime in which the magnitude of the outlier eigenvalue, which grows as 1*/*(|*ν*|^2^ − *g*^2^), is of order 1. In this regime, we find that *A* is partially aligned with eigenvectors of *J*, as described below. To obtain this result, we show that

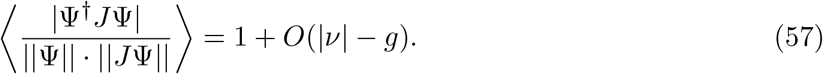

Note that the right-hand side being equal to 1 would imply that Ψ is an exact eigenvector of *J*, while a value of 0 is what we would expect from a random vector which is completely uncorrelated with *J*. Intermediate values correspond to partial alignment with eigenvectors of *J*.

In the large *N* limit, the norms (and their squared values) concentrate around their averages. Since we consider the limit from above, |*ν*| *> g*, these can be calculated directly by expanding the resolvent matrices as power series:

**Supplementary Figure 9.**
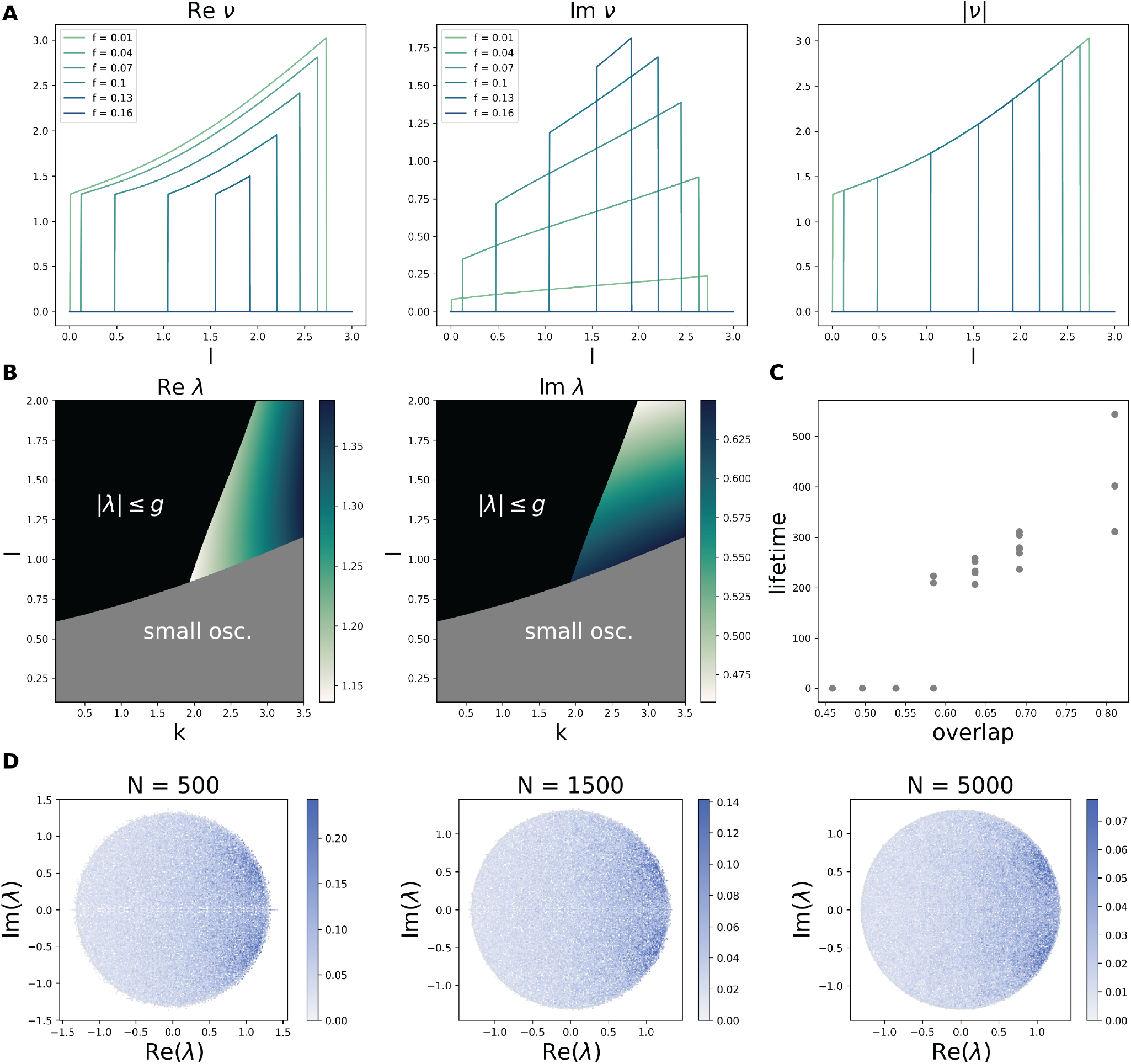
(A) Real and imaginary parts and modulus of *ν* as a function of input amplitude (*I*) and frequency (*f* ). The value of *ν* is set to zero whenever the eigenvalue outlier ceases to exist (|*λ*| ≤ *g*) or when the condition for the network to be inside the small oscillation regime holds (Eq. 46). Stronger inputs lead to a larger value of |*ν*|, and thus, smaller eigenvalue outliers (Eq. (55)). However, inputs which are excessively weak leave the network in the small oscillations regime, just as we find in simulations. Moreover, faster inputs lead to larger imaginary parts of *ν*. (B) Real and imaginary part of the eigenvalue outlier, *λ*, as a function of *k, I*, using the same parameter settings as in the main text Fig. 2. The outlier closely tracks the behavior of the lifetime. (C) Eigenvalue overlap calculated via Eq. (65) vs. median lifetime in simulations of the persistent oscillations, across networks with the same parameters. In the regime in which long-lasting persistent oscillations occur, the overlap is consistently close to 1. (D) Here we reproduce qualitatively the trend in Fig. 4Aiv at various values of *N* . Note that while individual overlap magnitudes decay as one would expect at large *N*, the matrix *A*(0) remains correlated with complex eigenvalue-eigenvector pairs. This is sufficient to create complex outliers.

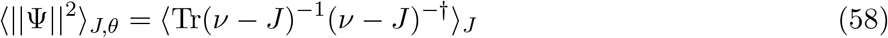

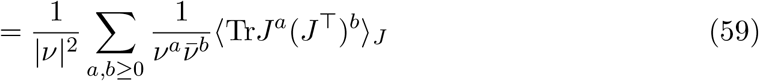

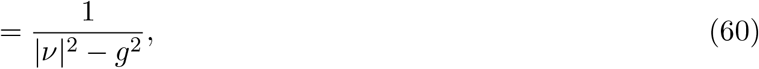

where we invoked the same argument used in the proof of Sec 2.3 to sum the series. Similarly:

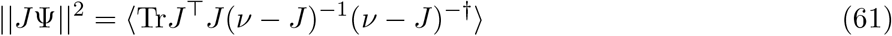

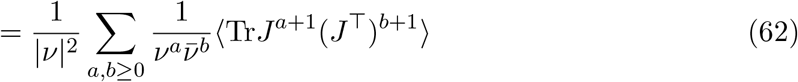

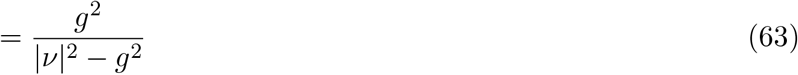

A similar line of reasoning shows that

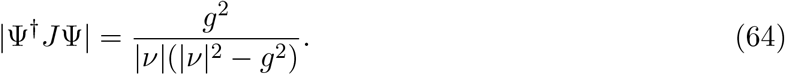

Thus:

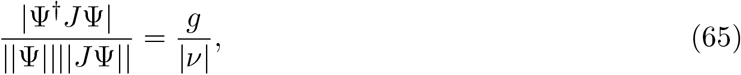

which gives the stated result after expanding in the limit |*ν*| − *g* → 0.

These results suggest that the eigenvalue outlier equations described in Secs. 2.2 and 2.3 collapse into the same formula in this limit. This is indeed the case. To see this, we first rescale the *α* appearing in the resolvent-aligned inputs of Sec. 2.3 as 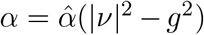 to compensate for the fact that the resolvent matrix, (*νI* − *J*)^−1^ diverges in the limit |*ν*| → *g*^+^. Writing *ν* = *ge*^*iφ*^ + *ϵ*, we can see that in the *ϵ* → 0^+^ limit, the outlier eigenvalues are given by

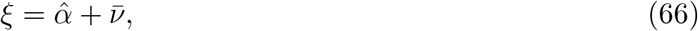

together with its conjugate, 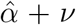. This result is what one would obtain from the eigenvector aligned perturbation of Sec. 2.2, demonstrating how these two results coincide as *ν* approaches the edge of the eigenvalue distribution.

## References

[1] H. Sompolinsky, A. Crisanti, and H.-J. Sommers, Chaos in random neural networks, Physical review letters 61, 259 (1988).

[2] J. Kadmon and H. Sompolinsky, Transition to chaos in random neuronal networks, Physical Review X 5, 041030 (2015).

[3] J. Aljadeff, M. Stern, and T. Sharpee, Transition to chaos in random networks with cell-type-specific connectivity, Physical review letters 114, 088101 (2015).

[4] F. Mastrogiuseppe and S. Ostojic, Linking connectivity, dynamics, and computations in low-rank recurrent neural networks, Neuron 99, 609 (2018).

[5] D. G. Clark, L. Abbott, and A. Litwin-Kumar, Dimension of activity in random neural networks, Physical Review Letters 131, 118401 (2023).

[6] J. J. Hopfield, Neural networks and physical systems with emergent collective computational abilities., Proceedings of the national academy of sciences 79, 2554 (1982).

[7] D. Krotov and J. Hopfield, Large associative memory problem in neurobiology and machine learning, arXiv preprint arXiv:2008.06996 (2020).

[8] H. Sompolinsky and I. Kanter, Temporal association in asymmetric neural networks, Physical review letters 57, 2861 (1986).

[9] M. Farrell and C. Pehlevan, Recall tempo of hebbian sequences depends on the interplay of hebbian kernel with tutor signal timing, Proceedings of the National Academy of Sciences 121, e2309876121 (2024).

[10] L. F. Abbott, J. Varela, K. Sen, and S. Nelson, Synaptic depression and cortical gain control, Science 275, 221 (1997).

[11] O. Barak and M. Tsodyks, Persistent activity in neural networks with dynamic synapses, PLoS computational biology 3, e35 (2007).

[12] G. Mongillo, O. Barak, and M. Tsodyks, Synaptic theory of working memory, Science 319, 1543 (2008).

[13] Y. Mi, M. Katkov, and M. Tsodyks, Synaptic correlates of working memory capacity, Neuron 93, 323 (2017).

[14] D. V. Buonomano and W. Maass, State-dependent computations: spatiotemporal processing in cortical networks, Nature Reviews Neuroscience 10, 113 (2009).

[15] A. Lansner, F. Fiebig, and P. Herman, Fast hebbian plasticity and working memory, Current Opinion in Neurobiology 83, 102809 (2023).

[16] R. C. Malenka and R. A. Nicoll, Nmda-receptor-dependent synaptic plasticity: multiple forms and mechanisms, Trends in neurosciences 16, 521 (1993).

[17] R. C. Malenka, Postsynaptic factors control the duration of synaptic enhancement in area ca1 of the hippocampus, Neuron 6, 53 (1991).

[18] B. Gustafsson, F. Asztely, E. Hanse, and H. Wigström, Onset characteristics of long-term potentiation in the guinea-pig hippocampal ca1 region in vitro, European Journal of Neuroscience 1, 382 (1989).

[19] M. A. Erickson, L. A. Maramara, and J. Lisman, A single brief burst induces glur1-dependent associative shortterm potentiation: a potential mechanism for short-term memory, Journal of cognitive neuroscience 22, 2530 (2010).

[20] F. Fiebig and A. Lansner, A spiking working memory model based on hebbian short-term potentiation, Journal of Neuroscience 37, 83 (2017).

[21] S. G. Manohar, N. Zokaei, S. J. Fallon, T. P. Vogels, and M. Husain, Neural mechanisms of attending to items in working memory, Neuroscience & Biobehavioral Reviews 101, 1 (2019).

[22] T. Miconi, K. Stanley, and J. Clune, Differentiable plasticity: training plastic neural networks with backpropagation, in International Conference on Machine Learning (PMLR, 2018) pp. 3559–3568.

[23] A. Sandberg, J. Tegnér, and A. Lansner, A working memory model based on fast hebbian learning, Network: Computation in Neural Systems 14, 789 (2003).

[24] J. Ba, G. E. Hinton, V. Mnih, J. Z. Leibo, and C. Ionescu, Using fast weights to attend to the recent past, Advances in neural information processing systems 29 (2016).

[25] D. G. Clark and L. F. Abbott, Theory of coupled neuronal-synaptic dynamics, Physical Review X 14, 021001 (2024).

[26] F. Schönsberg, D. Giana, Y. Chopra, M. Diamond, and S. Goldt, Diverse perceptual biases emerge from hebbian plasticity in a recurrent neural network model, bioRxiv,2024 (2024).

[27] K. Rajan, L. Abbott, and H. Sompolinsky, Stimulusdependent suppression of chaos in recurrent neural networks, Physical Review E–Statistical, Nonlinear, and Soft Matter Physics 82, 011903 (2010).

[28] That persistent oscillations can occur even when the activity is fully oscillatory is likely because full entrainment does not imply that the recurrence plays no role. In particular, near this phase transition, the recurrence still plays a large role in determining the amplitude and phases of the units, so the neural activity expresses features driven by both the inputs and the recurrence, leading to persistent oscillations.

[29] T. Tao, Outliers in the spectrum of iid matrices with bounded rank perturbations, Probability Theory and Related Fields 155, 231 (2013).

[30] D. J. Amit and N. Brunel, Model of global spontaneous activity and local structured activity during delay periods in the cerebral cortex., Cerebral cortex (New York, NY: 1991) 7, 237 (1997).

[31] X.-J. Wang, Synaptic reverberation underlying mnemonic persistent activity, Trends in neurosciences 24, 455 (2001).

[32] X.-J. Wang, 50 years of mnemonic persistent activity: quo vadis?, Trends in Neurosciences 44, 888 (2021).

[33] A. Compte, N. Brunel, P. S. Goldman-Rakic, and X.-J. Wang, Synaptic mechanisms and network dynamics underlying spatial working memory in a cortical network model, Cerebral cortex 10, 910 (2000).

[34] D. Mendoza-Halliday, S. Torres, and J. C. Martinez-Trujillo, Sharp emergence of feature-selective sustained activity along the dorsal visual pathway, Nature neuroscience 17, 1255 (2014).

[35] D. Zaksas and T. Pasternak, Directional signals in the prefrontal cortex and in area mt during a working memory for visual motion task, Journal of Neuroscience 26, 11726 (2006).

[36] B. Pesaran, J. S. Pezaris, M. Sahani, P. P. Mitra, and R. A. Andersen, Temporal structure in neuronal activity during working memory in macaque parietal cortex, Nature neuroscience 5, 805 (2002).

[37] E. Baeg, Y. Kim, K. Huh, I. Mook-Jung, H. Kim, and M. Jung, Dynamics of population code for working memory in the prefrontal cortex, Neuron 40, 177 (2003).

[38] J. Zhu, Q. Cheng, Y. Chen, H. Fan, Z. Han, R. Hou, Z. Chen, and C. T. Li, Transient delay-period activity of agranular insular cortex controls working memory maintenance in learning novel tasks, Neuron 105, 934 (2020).

[39] P. T. Sadtler, K. M. Quick, M. D. Golub, S. M. Chase, S. I. Ryu, E. C. Tyler-Kabara, B. M. Yu, and A. P. Batista, Neural constraints on learning, Nature 512, 423 (2014).

[40] M. D. Golub, P. T. Sadtler, E. R. Oby, K. M. Quick, S. I. Ryu, E. C. Tyler-Kabara, A. P. Batista, S. M. Chase, and B. M. Yu, Learning by neural reassociation, Nature neuroscience 21, 607 (2018).

[41] G. Dragoi and S. Tonegawa, Preplay of future place cell sequences by hippocampal cellular assemblies, Nature 469, 397 (2011).

[42] S. McKenzie, R. Huszár, D. F. English, K. Kim, F. Christensen, E. Yoon, and G. Buzsáki, Preexisting hippocampal network dynamics constrain optogenetically induced place fields, Neuron 109, 1040 (2021).

[43] A. Luczak, P. Barthó, and K. D. Harris, Spontaneous events outline the realm of possible sensory responses in neocortical populations, Neuron 62, 413 (2009).

[44] E. H. Park, S. Keeley, C. Savin, J. B. Ranck, and A. A. Fenton, How the internally organized direction sense is used to navigate, Neuron 101, 285 (2019).

[45] A. A. Fenton, Remapping revisited: how the hippocampus represents different spaces, Nature Reviews Neuro-science 25, 428 (2024).

[46] K. Duecker, K. B. Doelling, A. Breska, E. B. Coffey, D. V. Sivarao, and B. Zoefel, Challenges and approaches in the study of neural entrainment, Journal of Neuroscience 44 (2024).

[47] G. Sumbre, A. Muto, H. Baier, and M.-m. Poo, Entrained rhythmic activities of neuronal ensembles as perceptual memory of time interval, Nature 456, 102 (2008).

[48] J. P. Lerousseau, A. Trébuchon, B. Morillon, and D. Schön, Frequency selectivity of persistent cortical oscillatory responses to auditory rhythmic stimulation, Journal of Neuroscience 41, 7991 (2021).

[49] S. Van Bree, E. Sohoglu, M. H. Davis, and B. Zoefel, Sustained neural rhythms reveal endogenous oscillations supporting speech perception, PLoS biology 19, e3001142 (2021).

[50] L. T. Blanpain, E. R. Cole, E. Chen, J. K. Park, M. Y. Walelign, R. E. Gross, B. T. Cabaniss, J. T. Willie, and A. C. Singer, Multisensory flicker modulates widespread brain networks and reduces interictal epileptiform discharges, Nature communications 15, 3156 (2024).

[51] H. Lee, G. V. Simpson, N. K. Logothetis, and G. Rainer, Phase locking of single neuron activity to theta oscillations during working memory in monkey extrastriate visual cortex, Neuron 45, 147 (2005).

[52] J. Kamiński, A. Brzezicka, A. N. Mamelak, and U. Rutishauser, Combined phase-rate coding by persistently active neurons as a mechanism for maintaining multiple items in working memory in humans, Neuron 106, 256 (2020).

[53] M. Siegel, M. R. Warden, and E. K. Miller, Phasedependent neuronal coding of objects in short-term memory, Proceedings of the National Academy of Sciences 106, 21341 (2009).

[54] J. C. Magee and C. Grienberger, Synaptic plasticity forms and functions, Annual review of neuroscience 43, 95 (2020).

[55] P. Kidger, On Neural Differential Equations, Ph.D. thesis, University of Oxford (2021).

[56] A. J. Wakhloo, awakhloo/dynamic-memories: zen, 10.5281/zenodo.21499172 (2026).

## References

[1] Patrick Kidger. On Neural Differential Equations. PhD thesis, University of Oxford, 2021.

[2] John T Chalker and Bernhard Mehlig. Eigenvector statistics in non-hermitian random matrix ensembles. Physical review letters, 81(16):3367, 1998.

[3] Bernhard Mehlig and John T Chalker. Statistical properties of eigenvectors in non-hermitian gaussian random matrix ensembles. Journal of Mathematical Physics, 41(5):3233–3256, 2000.

[4] Terence Tao. Outliers in the spectrum of iid matrices with bounded rank perturbations. Probability Theory and Related Fields, 155(1):231–263, 2013.

[5] Joshua Feinberg and A. Zee. Non-hermitian random matrix theory: Method of hermitian reduction. Nuclear Physics B, 504(3):579–608, November 1997.

[6] Daniel Bessis, Claude Itzykson, and Jean-Bernard Zuber. Quantum field theory techniques in graphical enumeration. Advances in Applied Mathematics, 1(2):109–157, 1980.

[7] Joshua Feinberg. Non-hermitian random matrix theory: summation of planar diagrams, the ‘single-ring’theorem and the disc–annulus phase transition. Journal of Physics A: Mathematical and General, 39(32):10029, 2006.

